# Differential timing for glucose assimilation in *Prochlorococcus* and coexistent microbial populations at the North Pacific Subtropical Gyre

**DOI:** 10.1101/2021.10.04.462702

**Authors:** María del Carmen Muñoz-Marín, Solange Duhamel, Karin M. Björkman, Jonathan D. Magasin, Jesús Díez, David M. Karl, José M. García-Fernández

## Abstract

The marine cyanobacterium *Prochlorococcus* can utilize glucose as a source of carbon. However, the relative importance of inorganic and organic carbon assimilation and the timing of glucose assimilation are still poorly understood in these numerically dominant cyanobacteria. Here we investigated whole microbial community and group-specific primary production and glucose assimilation, using incubations with radioisotopes combined with flow cytometry cell sorting. We also studied changes in the microbial community structure in response to glucose enrichments and analyzed the transcription of *Prochlorocccus* genes involved in carbon metabolism and photosynthesis.

Our results showed a circadian rhythm for glucose assimilation in *Prochlorococcus*, with maximum assimilation during the midday and minimum at midnight, which was different compared with that of the total microbial community. This suggests that rhythms in glucose assimilation have been adapted in *Prochlorococcus* to couple the active transport to photosynthetic light reactions producing energy, and possibly to avoid competition from the rest of the microbial community. High-light *Prochlorococcus* strains showed most transcriptional changes upon glucose enrichment. Pathways involved in glucose metabolism as the pentose phosphate, the Entner-Dudoroff, glycolysis, respiration and glucose transport showed an increase in the transcript level. A few genes of the low-light strains showed opposite changes, suggesting that glucose assimilation has been subjected to diversification along the *Prochlorococcus* evolution.

## Introduction

*Prochlorococcus* is the most abundant photosynthetic organism on Earth, contributing to an important part of the total primary production (1–4). The outstanding relevance of this microorganism in the field of marine microbiology and ecology has been demonstrated by a large series of studies published since its discovery, ca. 35 years ago (5). Because of its abundance, *Prochlorococcus* is also one of the main microbial players in biogeochemical cycles (6).

Early studies on this cyanobacterium were focused on photosynthesis, and it was widely considered an obligate photolithoautotrophic organism. However, it has been showed recently that *Prochlorococcus* can take up and use organic compounds, such as amino acids (containing nitrogen) (7, 8), DMSP (containing sulfur) (9) or phosphonates and adenosine triphosphate (ATP) (containing phosphorus) (10, 11), which recently has been reviewed (12). Since *Prochlorococcus* is adapted to thrive in very oligotrophic regions of the ocean, the use of those organic molecules was thought to be linked to their content in limiting elements. In this context, the discovery of glucose assimilation in *Prochlorococcus* was surprising (13), since this molecule is devoid of limiting element, containing only carbon, oxygen and hydrogen. However, it also contains potential energy which could be used by *Prochlorococcus*. Glucose addition to *Prochlorococcus* culture medium induced changes in the expression of a number of genes, including *glcH*, which encodes a multiphasic transporter with high affinity constant (Ks) in the nanomolar range (14). The ubiquity of this gene in all sequenced genomes of *Prochlorococcus* and marine *Synechococcus*, and the diversity of kinetics of the transporter (K_s_ and V_max_ parameters) (15), suggest that *glcH* is very important for *Prochlorococcus* and *Synechococcus*, and has been subjected to selective evolution in their genomes (12, 15).

The effects of glucose addition to *Prochlorococcus* in culture isolates showed specific increases in the expression of genes related to glucose metabolism (13). Proteomic analysis showed some changes which were reproducible but quantitatively small (15), including in proteins related to glucose metabolism. Besides, the expression of *glcH* has been shown to increase with higher concentrations of glucose in *Prochlorococcus* cultures, but to decrease in the dark (16). However, less is known about *Prochlorococcus* glucose utilization in the wild: glucose assimilation was demonstrated in natural populations of *Prochlorococcus* in the Atlantic (14) and in the Southwest Pacific (11) oceans; furthermore, it was shown that *Prochlorococcus* glucose assimilation in the field was reduced in the dark and in the presence of photosynthesis inhibitors (11).

Here, we further address *Prochlorococcus* glucose metabolism in field experiments carried out at Station ALOHA in the North Pacific Subtropical Gyre (NPSG) to determine how glucose addition affects the total microbial community during light-dark cycles. We measured glucose turnover rates and assimilation by the whole microbial community as well as in flow sorted *Prochlorococcus* over a diel cycle. Paired experiments measuring primary production were conducted in order to assess the relative contribution of glucose assimilation to *Prochlorococcus* total carbon assimilation. We used metagenomics to investigate the effect of glucose enrichment on the composition of natural populations (both picocyanobacteria and heterotrophic bacteria) and analyzed *Prochlorococcus* populations using microarrays to identify possible changes in the transcription of genes involved in carbon metabolism and photosynthesis pathways.

## Material and Methods

### Field sampling

Seawater for all incubation experiments was collected at Station ALOHA (22° 45’ N, 158° 00’ W) in the NPSG, during the KM1715 (HOT-296) cruise on October 5–9^th^, 2017. The seawater was sampled from a depth of 7 m using the RV Kilo Moana’s uncontaminated seawater system (https://www.soest.hawaii.edu/UMC/cms/KiloMoana.php). The surface light flux values measured during the experiments are shown in Table S1.

### Incubation experiments with radiolabeled substrates

Separate incubation experiments were carried out to determine primary production, using ^14^C-sodium bicarbonate (MP Biomedical 117441H; specific activity 2.22 TBq mmol^−1^), and the turnover and assimilation of glucose by *Prochlorococcus* and the whole microbial community, using both ^14^C-glucose and ^3^H-glucose (NEC042X (^14^C (U)); specific activity 9.7 GBq mmol^−1^ and 97% radiochemical purity, Perkin-Elmer NET100C (D-[6-^3^H(N)]); specific activity 1.83 TBq mmol^−1^ and 97% radiochemical purity). The radiolabeled glucose was also used to estimate the ambient glucose concentration in the seawater. These incubations were performed in 60 ml or 100 ml clear polycarbonate bottles that had been acid cleaned and ultra-pure water rinsed before rinsing with sample seawater. The seawater samples were spiked with the relevant radioisotope and incubated in on deck incubators with surface seawater cooling and blue shielding at 60% of full surface light.

### Estimation of ambient glucose concentrations

To estimate the ambient concentration of glucose in the surface seawaters at Station ALOHA, concentration series bioassays were carried out on the HOT 295, 296 and 298 cruises (Sep, Oct and Dec 2017, respectively) following the procedure described by Wright and Hobbie (17, 18) and modified by Zubkov and Tarran (19). Surface seawater was amended with increasing amounts of radiolabeled glucose, either as ^3^H-glucose (HOT 296, 298) or ^14^C-glucose (HOT 295, 298), incubated in the on-deck incubators and subsampled in a time-course fashion. For that, radioactive glucose was added at five to six different concentrations within a target range of 0.2–2.0 nmol glucose l^−1^. Each target concentration was run as duplicate incubations. The incubations were subsampled after 30, 60, 90, 120 and 240 minutes. The 4 hours timepoint was used as validation for the 4 hours incubation times used in the diel study. On HOT 295 incubations started at 12:30 h and used ^14^C-glucose only. During HOT 296 two bioassays were run using ^3^H-glucose with incubations started at 07:10 h on 6^th^ Oct, and at 09:40 on the 7^th^ of Oct. Two additional bioassays were performed on HOT 298 using both ^3^H- and ^14^C-glucose and started at noon Dec 14^th^. At each sampling time, 10 ml were filtered onto 0.2 μm polycarbonate filters, rinsed with filtered seawater and placed into plastic scintillation vials (Simport snap-twist vials, 7 ml). A small subsample (25 μl) was also collected from each incubation bottle to determine the total radioactivity used to calculate the added glucose concentration from the specific activity provided by the manufacturer of each isotope.

Glucose turnover times were determined by dividing the total radioactivity by the assimilation rate, where the rate of assimilation was derived from the linear regression of the increase in particulate radioactivity over time. By plotting the turnover times against the concentrations of added glucose, the ambient concentration of glucose can be derived from where the linear regression line intersects the x-axis (y=0, absolute value). The final glucose assimilation rates were calculated from the ambient + added glucose concentrations. Further details of the bioassay method have been described by Zubkov et al. (20).

### Diel assimilation

Seawater samples were collected every 6 h for a total of 10 samplings over a 54-h period. Duplicate 60 ml bottles were spiked with radiolabeled glucose to a target addition of 2 nM glucose or with ^14^C-sodium bicarbonate (final activity approximately 150 MBq l^−1^). Each sample set included a paraformaldehyde killed control (0.24 % final concentration) that was incubated alongside the live samples. The diel samples were incubated for 4 h and terminated by adding paraformaldehyde (0.24% final concentration) to stop the assimilation of the radiolabel and preserve the sample. The primary production incubations were treated the same way as the glucose incubations during daylight hours but these samples were kept in the incubator overnight (from 18:30 to 04:00), under the assumption that primary productivity during the night would be negligible. At the end of the incubation period, 5 ml subsamples from each incubation were filtered onto 0.2 μm polycarbonate filters, rinsed with filtered seawater and placed either into plastic scintillation vials (Simport snap-twist scintillation vials, 7 ml) for glucose or into 20 ml borosilicate scintillation vials for ^14^C-sodium bicarbonate. The latter were acidified (1 ml 2N HCl) and vented for 24 h before the addition of scintillation cocktail. The total radioactivity subsamples for ^14^C-bicarbonate were trapped with phenethylamine (Sigma Aldrich 407267). Ultima Gold LLT (Perkin-Elmer) was used as the scintillation cocktail for all radioassays in a Packard Tri-Carb® liquid scintillation counter. Dissolved inorganic carbon concentrations for HOT 296 were from the HOT Data Organization and Graphical System (http://hahana.soest.hawaii.edu/hot/hot-dogs), and a correction of 1.06 for preferential assimilation of ^12^C relative to ^14^C was applied (21).

Subsamples for flow cytometric cell counts were taken in duplicate 2 ml samples, fixed with paraformaldehyde (0.24% final concentration) and incubated for 15 min in the dark before flash freezing and storing at −80°C until analysis back in the laboratory. The remaining sample volume from each incubation was concentrated as described in Duhamel et al. (3, 11) for cell sorting to determine cell and group specific assimilation of glucose and inorganic carbon by *Prochlorococcus*.

### Flow cytometry counting and cell sorting

Phytoplankton groups were enumerated using a BD Influx flow cytometer equipped with a forward scatter (FSC) detector with small particle option (BD Biosciences, San Jose, CA, USA). *Prochlorococcus*, *Synechococcus, Crocosphaera* and pigmented eukaryotes (<5-μm (3)) were enumerated in unstained samples following published protocols (22). Briefly, cells were identified based on red fluorescence signals vs FSC, then further gated by FSC and orange fluorescence (Fig. S1). The high phycoerythrin (orange) signal in *Synechococcus* and *Crocosphaera* was used to distinguish them from *Prochlorococcus* and pigmented eukaryotes. A 488 plus a 457 nm (200 and 300 mW solid state, respectively) laser focused into the same pinhole resolved dim surface *Prochlorococcus* population from background noise in a FSC vs red fluorescence plot. Potential particle aggregates were discarded using a pulse width vs. forward scatter plot. Calibration and alignment were done using 1-μm yellow-green microspheres (Polysciences, USA).

Group-specific rates of ^3^H-glucose /^14^C-glucose assimilation and primary production by *Prochlorococcus* were determined by measuring the amount of radioactivity assimilated into populations sorted using the BD Influx (100 μm nozzle tip, sheath solution (sodium chloride 6 g L^−1^ in ultrapure water and filtered in-line through a 0.22-μm Sterivex™ filter unit), 1.0 drop single mode) according to Duhamel et al., (3, 11).

The drop delay was calibrated using Accudrop Beads (BD Biosciences, USA) and sorting efficiency was verified manually by sorting a specified number of 1-μm yellow-green microspheres (Polysciences, USA) onto a glass slide and counting the beads under an epifluorescence microscope. We systematically recovered 100% of the targeted beads before sorting cells. For each sample, 50,000 *Prochlorococcus* cells were sorted, filtered onto 0.2-μm polycarbonate membranes, rinsed with filtered seawater, and assayed by liquid scintillation counting (dpm cell^−1^). The ^14^C-labeled samples were acidified with 1 mL of 2M HCl for 24 h to remove any unincorporated ^14^C-sodium bicarbonate before adding the scintillation cocktail. Radioactivity per cell (dpm cell^−1^) measured in the killed control samples was subtracted from radioactivity per cell measured in the respective sample. On average, radioactivity in the killed controls for 50,000 *Prochlorococcus* cells sorted (i.e., blanks) was 5 ± 2×10^−5^ and 4.7 ± 1.3×10^−4^ dpm cell^−1^ for ^3^H-glucose and ^14^C-sodium bicarbonate, respectively. Detection limits are defined as 2X the killed control before it being subtracted from the sample. The cell-specific assimilation rate (nmol cell^−1^ h^−1^) was calculated by dividing the radioactivity per cell (dpm cell^−1^) by the total microbial activity (dpm l^−1^) measured in the same treatment, and then multiplied by the total microbial assimilation rate at ambient plus added organic substrate concentration (S_a_+S*, nmol l^−1^ h^−1^) as described in Duhamel and coworkers (11).

### Incubation experiments with glucose for genomics-based approaches

For molecular approaches seawater was collected into 2 L polycarbonate bottles in triplicate control and glucose amended samples. The glucose treatments were spiked with 0.1 μM of non-radiolabeled glucose and incubated alongside the controls (unamended) in the on-deck incubators. For the first experiment (October 6^th^), surface seawater was sampled at noon, while for the second experiment (October 7^th^) surface seawater was collected at 16:00.

Samples were collected for these two experiments, and for each one, two technical replicates were collected at 3 different times: after 4h, 12h and 24h of incubation, in the presence of glucose or in the control treatment. Therefore, samples are identified as follows: 4h_glucose, 4h_control, 12h_glucose, 12h_control, 24h_glucose, 24h_control, 2D_4h_glucose, 2D_4h_control, 2D_12h_glucose, 2D_12h_control, 2D_24h_glucose, 2D_24h_control with the second experiment identified as “2D” and considered as biological replicates. The different technical replicates are labeled with the number 1 or 2 before the treatment, e.g. (4h_1glucose and 4h_2glucose.

At each sampling time point, RNA and DNA samples were collected by filtering 4 L and 1 L of seawater, respectively onto 0.22 μm pore-size Sterivex cartridges (Millipore Corp., Billerica, MA, USA) using a peristaltic pump set at low rate to maintain low pressure. Filters were opened and carefully placed in sterile 2 ml bead-beating tubes with sterile glass beads and stored at −80 °C until extraction.

### DNA extraction

DNA extractions were carried out with a modification of the Qiagen DNeasy Plant Kit (23). Briefly, 400 μl lysis buffer (AP1 buffer) was added to the bead-beating tubes, followed by three sequential freeze-thaw cycles using liquid nitrogen and a 65°C water bath. The tubes were agitated for 2 min with a Vortex-Genie 2 bead beater (Scientific Industries, Inc), and incubated for 1 h at 55°C with 20 mg ml^−1^ proteinase K (Qiagen). Samples were treated for 10 min at 65 °C with 4 μl RNase A (100 mg ml^−1^) and then the filters were removed using sterile needles. The tubes were centrifuged for 5 min at 16,873 x g at 4°C, and the supernatant was further purified using the manufacturer’s protocol (Qiagen). Samples were eluted using 100 μl of the elution buffer (AE buffer) and stored at 20°C.

Sufficient environmental DNA was obtained for two technical replicates only in 5 samples (2D_4h_glucose, 2D_12h_glucose, 2D_12h_control, 2D_24h_glucose, 2D_24h_glucose) and those samples are identified as follows: 2D_4h_1glucose, 2D_4h_2glucose, 2D_12h_1glucose, 2D_12h_2glucose, 2D_12h_1control, 2D_12h_2control, 2D_24h_1glucose, 2D_24h_2glucose, 2D_24h_1glucose and 2D_24h_2glucose.

### Sequence processing

V3 and V4 regions of 16S rRNA genes were amplified, sequenced and analyzed by the STAB-VIDA company (Lisbon, Portugal) using the following primers (24): forward primer 5’CCTACGGGNGGCWGCAG-3 and reverse primer 5’-GACTACHVGGGTATCTAATCC-3.

DNA samples were checked for quantity and integrity by 1% Agarose gel electrophoresis, a Qubit® Fluorometer (Thermo Fisher Scientific, MA, USA) and a 2100 Bioanalyzer (Agilent Technologies, Santa Clara, CA, USA) prior to library construction using the Illumina 16S Metagenomic Sequencing Library Preparation protocol (25). The generated DNA fragments (DNA libraries) were sequenced with the lllumina MiSeq platform using MiSeq Reagent Kit v2 to produce paired-end sequencing reads (2×250 bp). FastQC (26) was used to inspect the quality of the raw sequencing reads. The analysis of the generated raw sequence data was carried out using QIIME2 v2018.2 (27). The QIIME2 plugin for DADA2 (denoise-paired) (28) was used to process the raw reads into amplicon sequence variants (ASVs) which provide higher phylogenetic resolution compared to operational taxonomic units (OTUs). Reads were trimmed of primers at the 5’ end using the primer lengths and truncated at the 3’ end so that total lengths were 250 bp (R1) and 235 bp (R2). Reads were removed if they had Phred quality scores <20 on average, or <17 for two consecutive bases, or if they had >2 expected errors. Quality-filtered reads were then dereplicated, denoised (ASV inference using the core DADA2 algorithm), merged, and filtered for chimeras.

The 1534 ASVs identified by DADA2 were additionally filtered for chimera using the uchime3_denovo algorithm implemented in vsearch v2.13.3 (29) which identified 53 chimera, and the NCBI 16S rRNA chimera detection pipeline based on uchime2_ref (30) which identified 452 chimera. One of the 452 chimera (ASV.348) was retained because it was observed in 12 samples (198 sequences total) and had taxonomically consistent parents (genus *Coxiella*). From the 1030 non-chimeric ASVs, we removed 231 ASVs with ≤10 total sequences and another 81 ASVs that were detected in only 1 sample with <30 sequences, to produce a final set of 718 ASVs (Table S2). The ASVs were classified by taxon using a QIIME2 scikit-learn fitted classifier that had been trained on the SILVA database (release 128 QIIME) clustered at 97% similarity (Table S2).

### RNA extraction and processing for hybridization to the microarray

Environmental RNA containing transcripts from *Prochlorococcus* cells was extracted using an Ambion RiboPure Bacteria kit (Ambion, Thermo Fisher), with modifications that included mechanical lysis using glass beads (Biospec, Bartlesville, OK). The extracted RNA was treated with a Turbo-DNA-free DNase kit (Ambion, Thermo Fisher) to remove genomic DNA. Samples were collected for two experiments, for each one two technical replicates were collected at 3 different time points, same as DNA samples. Sufficient environmental RNA was obtained for two technical replicates in 4 samplings (4h_1control, 4h_2control, 4h_1glucose, 4h_2glucose, 12h_1glucose, 12h_2glucose, 2D_4h_1glucose and 2D_4h_2glucose).

RNA concentration, purity and quality were determined using a NanoDrop 1000 instrument (Thermo Scientific, Waltham, MA, USA), a 2100 Bioanalyzer (Agilent Technologies, Santa Clara, CA, USA), and an RNA 6000 Nano kit (Agilent Technologies). Only samples with RNA integrity values of >7.0 and ratios of A_260_/A_230_ and A_260_/A_280_ ≥1.8 were processed further. Double-stranded cDNA (ds-cDNA) was synthesized from environmental RNA samples that contained *Prochlorococcus* and amplified following the procedure previously described by Shilova et al. (31). Briefly, 400 ng RNA from each sample was used, and 1 μl of a 1:100 dilution (corresponding to 4.7 aM of ERCC-0016) of RNA spike-in mix 1 (External RNA Control Consortium (32) (Ambion)) was added before amplification was performed to monitor the technical performance of the assay showing linear amplification of specific probes (Fig. S2) (32). Double-stranded cDNA was synthesized and amplified using a TransPlex whole-transcriptome amplification kit (WTA-2; Sigma-Aldrich, St. Louis, MO, USA) and antibody-inactivated hot-start Taq DNA polymerase (Sigma-Aldrich). The amplified cDNA was purified with a GenElute PCR cleanup kit (Sigma-Aldrich), and concentration, purity and quality of ds-cDNA were determined using a NanoDrop 1000 instrument, a 2100 Bioanalyzer, and an Agilent DNA 7500 kit (Agilent Technologies). Total RNA concentration of 400 ng yielded on average 12 μg of ds-cDNA. The labeling and hybridization of cDNA samples (2.0 μg of ds-cDNA) to the microarray were done at the facility Centro de Investigación Principe Felipe (Valencia, Spain) according to the Agilent Technology protocol for arrays.

### Design of the Prochlorococcus array

The *Prochlorococcus* oligonucleotide expression array was designed using *Prochlorococcus* genes and the eArray Web-based tool (Agilent Technology Inc.; https://earray.chem.agilent.com/earray/) similarly to the array design previously described by (31, 33). The gene sequences were obtained from the National Center of Biotechnology Information (NCBI; https://www.ncbi.nlm.nih.gov). Briefly, six probes of 60 nucleotides in length were designed for each gene, and a total of 7,501 probes (1,326 genes) were designed for *Prochlorococcus*. The probes were designed based on the sequenced genomes of the strains more abundant at Station ALOHA for specific core genes involved in carbon metabolism and photosynthesis pathways. These probes were replicated 4 times in the 8 × 60K array slides, which allowed internal evaluation of signals. The sequences of all oligonucleotide probes were tested *in silico* for possible cross-hybridization as described below. The probe sequences were used as queries in the BLASTN against the following available nt databases in June 2017: Marine microbes, Microbial Eukaryote Transcription, and Non-redundant Nucleotides NCBI SRA website and all rRNA databases from Silva as of February 2, 2016.

Agilent technology allows 5% nt mismatch in the whole probe region; thus, sequences with a range of 95% to 100% nt identity to the target probe are detected. Therefore, all probes with BLASTN hits with ≥95% over 100% of the nt length were deleted. Next, the probe sequences that passed the cross-hybridization filter were clustered using CD-HIT-EST (34, 35) at 95% nt similarity to select unique probes for *Prochlorococcus*.

In addition, standard control probes (IS-62976-8-V2_60Kby8_GX_EQC_201000210 with ERCC control probes added) were included randomly as part of the Agilent Technology array to feature locations on the microarray slide. The final design of the microarray was synthesized on a platforms of ca. 62,976 experimental probes and 1,319 control probes on each 8 × 60K array slide. The probe sequences are available at NCBI Gene Expression Omnibus (GEO) under accession number GSE154594.

### Microarray data analysis

All data analyses were performed with R (https://www.R-project.org) and packages from the Bioconductor Project (36), specifically, using the Biobase (37), Linear Models for Microarray LIMMA (38), arrayQualityMetrics (39) and affyPLM (40, 41). These packages were mainly utilized via software that was developed for the MicroTOOLs environmental microarray (31, 42), which we adapted slightly to the *Prochlorococcus* microarrays. As in the prior study (42) arrays were normalized by quantiles and gene intensities were calculated by median polishing (Fig. S3). Gene detection was done separately for each gene in each sample. Specifically, each gene *g* was detected in each sample *s* if it had a signal to noise ratio *SNR*_*gs*_ ≥ *5*, where *SNR*_*gs*_ = *S*_*gi*_ / *BG*_*s*_ and *BG*_*s*_ was the background intensity in *s*. We defined *BG*_*s*_ based on the lowest detected ERCC mRNA spike-in transcripts. For each sample ERCC spike-in transcript intensities were linearly modeled (Fig. S2). Then we identified in *s* the least concentrated ERCC with a modelled intensity that was twice the median of measured intensities for Agilent negative control probes (structural hairpins). *BG*_*s*_ was the modelled intensity for this ERCC. On average 448 genes (mean) were detected in each sample (min 416, max 538). In total, 775 detected genes were detected across the samples (union). Raw and normalized microarray data for *Prochlorococcus* were submitted to NCBI GEO under accession number GSE154594.

As in the prior study (42), differentially expressed (DE) genes were identified using the LIMMA functions lmFit, eBayes, and topTreat. Empirical Bayes is well suited to studies with few samples because it pools them to estimate the variances for each gene’s linear model (43). To identify biologically relevant DE, we looked for genes with fold changes that were at least 1.3× different (not simply >0) between treatments and matched controls (Benjamini-Hochberg adjusted *p*-value < 0.05). DE genes always had changes >1.5-fold (mean 2.3-fold) and were mainly identified in the experiment 2 at 12h (2D_12h_glucose vs. 2D_12h_control). DE genes were required to be above detection cut offs in at least one of the treatment or control samples.

The Ensemble Gene Set Enrichment Analyses approach (EGSEA; (44)) was used to check for significant transcript level changes that occurred collectively for genes from the same pathway. Briefly, for each *Prochlorococcus* clade, genes were assigned to pathways based on the literature (with multiple pathway memberships allowed; Supplemental Information). Only genes for which transcripts tend to change in the same direction in a pathway (increasing or decreasing together) were included in our pathways.

## Results

### Prochlorococcus cell abundance and cell size

*Prochlorococcus* dominated phytoplankton abundances, with on average 1.23 ± 0.35 × 10^5^ cell ml^−1^, with *Synechococcus*, *Crocosphaera* and picophytoeukaryotes contributing 1.47 ± 0.39 × 10^3^, 3.77 ± 0.97 × 10^2^, 8.32 ± 1.92 × 10^2^ cell ml^−1^, respectively (average ± standard deviation, n = 20 (biological and technical replicates; Fig. S4).

*Prochlorococcus* cell abundance and cell size showed a diel cycle with increasing cell abundance and cell diameter during the daylight period and lowest values during the dark period. Cell abundances varied from 0.7 × 10^5^ to 1.8 × 10^5^ cell ml^−1^, while cell size varied from 0.36 to 0.41 μm (Fig. S5).

### Inorganic carbon fixation rates (primary production)

Rates of inorganic carbon fixation by the whole community (> 0.2 μm, Fig. 1a, Table 1) ranged from 0.8 ± 0.0 nmol C l^−1^ h^−1^ at night (18:30 to 4:00 incubations) to 28.6 ± 0.8 nmol C l^−1^ h^−1^ at noon (noon to 16:00 incubations). On average, carbon fixation during daylight was 25.1 ± 3.1 nmol C l^−1^ h^−1^ (n=12). On a per cell level, *Prochlorococcus* also showed a pronounced diel cycling in carbon fixation with undetectable values at night and 13.1 ± 8.5 nmol l^−1^ h^−1^ (1.22 ± 0.66 fg C cell^−1^ h^−1^) during the day (n = 12, Fig. 1b, Table 1). As a taxon specific group, *Prochlorococcus* represented 41.5 ± 16.5% of the total carbon fixation during the day (34.8 ± 10% in the morning and 49.6 ± 20% in the afternoon (n = 11)).

**Figure 1.**
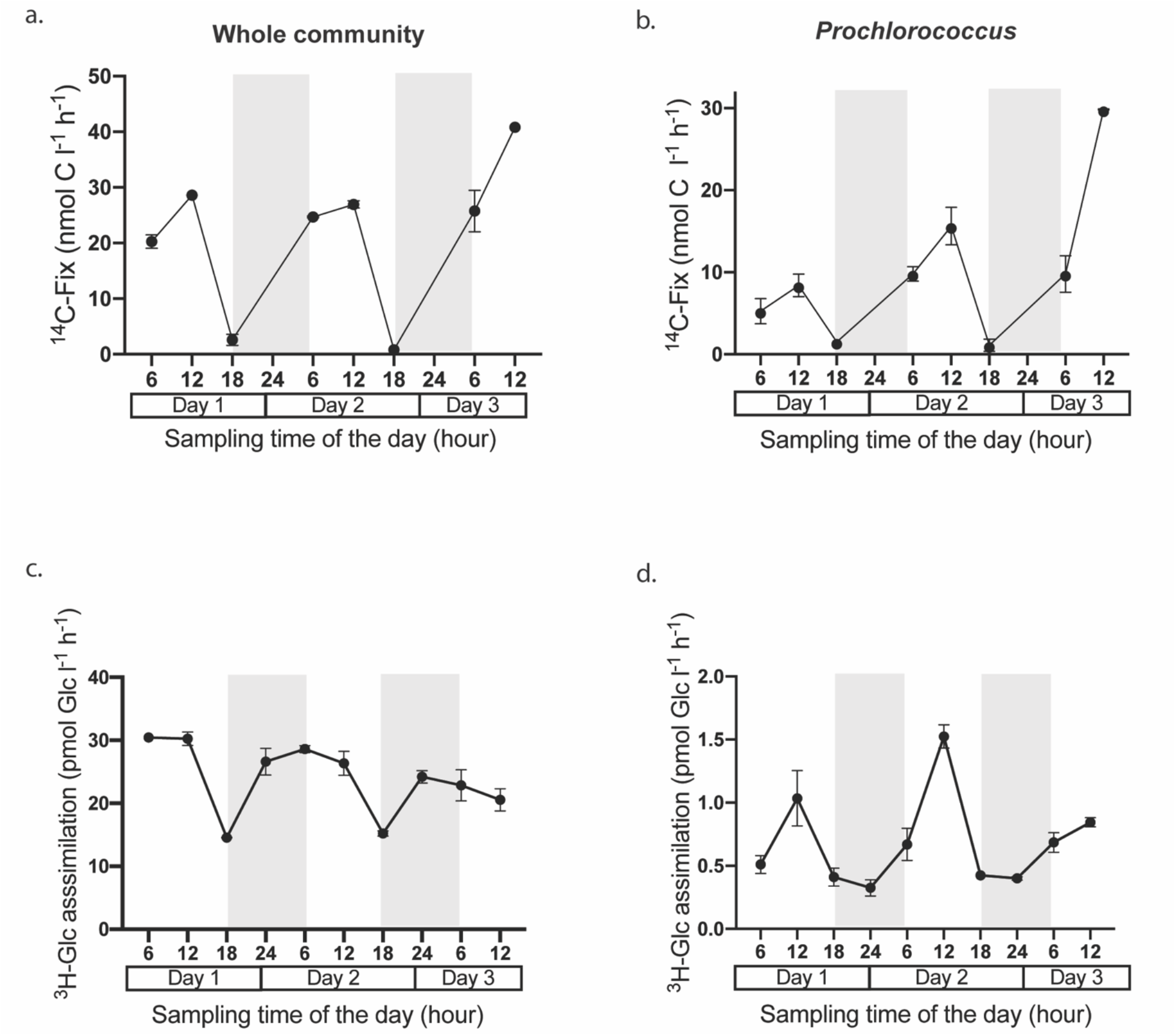
Top panel (a, b): Inorganic carbon fixation rates over time by the whole community (a; total, > 0.2 μm; nmol C l^−1^ h^−1^), and by *Prochlorococcus* (b; nmol C l^−1^ h^−1^). Bottom panel (c, d): Glucose assimilation over time by the whole community (c; total, > 0.2 μm; pmol Glc l^−1^ h^−1^), and by *Prochlorococcus* as a group (d; pmol Glc l^−1^ h^−1^). The shaded area represents the dark period.

**Table 1.** Rates of inorganic carbon fixation (^14^C-PP; primary productivity) and assimilation of glucose (^3^H-Glu and ^14^C-Glu) by the whole water communities (WW), *Prochlorococcus* cell abundance (PRO cell#), *Prochlorococcus* sodium bicarbonate fixation (^14^C-PP-PRO) and glucose assimilation (^3^H-Glc-PRO) during the diel study. The values presented in the table are average of two technical replicates and the standard deviation. Percentages (%) in bold in the ^14^C-PP-PRO column show the relative contribution by *Prochlorococcus* to total (WW) primary production. The ^3^H-Glc-PRO column shows the % glucose assimilation by *Prochlorococcus* relative to community glucose assimilation (WW), and the carbon contribution from glucose to inorganic carbon fixation by *Prochlorococcus* (Pro).

**Titles and legends to main figures**
| Sample # | Hour of the day | <sup>14</sup>C-PP-WW (nmol C l<sup>-1</sup> h<sup>-1</sup>) | <sup>3</sup>H-Glc-WW (pmol C l<sup>-1</sup> h<sup>-1</sup>) | <sup>14</sup>C-Glc-WW (pmol C l<sup>-1</sup> h<sup>-1</sup>) | PRO cell# (10<sup>8</sup> l<sup>-1</sup>) | <sup>14</sup>C-PP-PRO (nmol l<sup>-1</sup> h<sup>-1</sup>) | <sup>3</sup>H-Glc-PRO (pmol l<sup>-1</sup> h<sup>-1</sup>) |
| --- | --- | --- | --- | --- | --- | --- | --- |
| 1 (Day 1) | 6-10 | 20.3±1.2 | 30.4±0.3 | 64.5±13.2 | 0.89 | 5.25±1.54<br>**26% ww** | 0.51±0.06<br>**1.7 % ww**<br>**0.06% Pro** |
| 2 | 12-16 | 28.6±0.8 | 30.3±1.1 | 54.1±2 | 1.41 | 8.40±1.38<br>**30% ww** | 1.03±0.21<br>**3.4% ww**<br>**0.08% Pro** |
| 3 | 18-22 | 2.6±1.0 | 14.5±0.1 | 28.2±0.6 | 1.57 | 1.51±0.45<br>**67% ww** | 0.40±0.06<br>**2.8% ww**<br>**0.18% Pro** |
| 4 | 24-4 | - | 26.6±2.1 | 55.2 | 0.80 | - | 0.32±0.06<br>**1.23% ww** |
| 5 (Day 2) | 6-10 | 24.7±0.1 | 28.6±0.5 | 57.1±1.1 | 0.88 | 9.79±0.88<br>**40% ww** | 0.67±0.13<br>**2.3% ww**<br>**0.04% Pro** |
| 6 | 12-16 | 27.0±0.7 | 26.3±1.9 | 55.2±1.9 | 1.90 | 15.62±2.27<br>**58% ww** | 1.52±0.00<br>**5.8% ww**<br>**0.06% Pro** |
| 7 | 18-22 | 0.8±0.00 | 15.2±0.4 | 28.7±1.4 | 1.43 | 1.10±0.75<br>**100% ww** | 0.42±0.01<br>**2.8% ww**<br>**0.31% Pro** |
| 8 | 24-4 | - | 24.2±1.0 | 45.1±1.4 | 0.93 | - | 0.40±0.01<br>**1.65% ww** |
| 9 (Day 3) | 6-10 | 25.7±3.7 | 23.0±2.4 | 46.0±10.2 | 1.10 | 9.79±2.22<br>**39% ww** | 0.69±0.08<br>**3% ww**<br>**0.04% Pro** |
| 10 | 12-16 | 23.8 | 20.6±1.7 | 41.4±1.9 | 1.40 | 29.83±0.06<br>**100% ww** | 0.85±0.04<br>**4.1% ww**<br>**0.02% Pro** |

### Glucose assimilation

The two bioassays conducted using ^3^H-glucose during HOT 296 indicated an ambient glucose concentration of 1.1 ± 0.1 nmol l^−1^. The ^3^H-glucose spike added, on average, an additional 1.9 ± 0.2 nmol l^−1^ (n=20). Taking both the ambient and added glucose concentrations into account, the assimilation of glucose by the whole community (> 0.2 μm, Fig. 1c, Table 1) varied over the diel cycle by approximately a factor of 2 (14.5 ± 0.1 to 30.4 ± 0.3 pmol ^3^H-Glc l^−1^ h^−1^), with on average 24.0 ± 5.6 pmol ^3^H-Glc l^−1^ h^−1^ (n = 20). A diel pattern was observed for the whole community with lower values in incubations started at dusk (18:00), averaging 14.9 ± 0.5 pmol ^3^H-Glc l^−1^ h^−1^ (n = 4) and higher values in incubations started at midnight and early morning averaging 25.6 ± 2.7 pmol ^3^H-Glc l^−1^ h^−1^ (n = 8) (Fig. 1c, Table 1).

However, *Prochlorococcus* showed a different diel pattern in glucose assimilation with 0.003 ± 0.001 amol Glc cell^−1^ h^−1^ at night (n = 8) and 0.007 ± 0.001 amol Glc cell^−1^ h^−1^ during the day (n = 12, Table 2). Similarly, as a taxon specific group, *Prochlorococcus* presented a pronounced diel cycle with maximum values at noon of 0.88 ± 0.36 pmol Glc l^−1^ h^−1^ (n=6) and minimum values at midnight of 0.39 ± 0.05 pmol Glc l^−1^ h^−1^ (approximately 2.3-fold change) (n = 4, Fig. 1d, Table 1).

**Table 2.** Rates of inorganic carbon fixation and glucose assimilation by the whole community, and by *Prochlorococcus* and *Synechococcus* reported for natural samples. In this study, the detection limits are defined as 2X the blank before it being subtracted from the sample.

|  | *Whole community* | *Prochlorococcus* | *Synechococcus* |
| --- | --- | --- | --- |
| **Carbon fixation**<br>(<sup>14</sup>C-sodium bicarbonate) | 26.4 day/<br>undetectable at night<br>(nmol l<sup>-1</sup> h<sup>-1</sup>)<sup>a</sup> | 1.22 day / undetectable at night (fg C cell<sup>-1</sup> h<sup>-1</sup>)<sup>a</sup><br>or 13.1 day/ undetectable at night (nmol l<sup>-1</sup> h<sup>-1</sup>)<sup>a*</sup><br>0.9 to 1.6 (fg C cell<sup>-1</sup> h<sup>-1</sup>)<sup>b</sup><br>1.2 (fg C cell<sup>-1</sup> h<sup>-1</sup>)<sup>c</sup> |  |
| **Glucose**<br>(<sup>3</sup>H-glucose) | 24<br>(pmol l<sup>-1</sup> h<sup>-1</sup>)<sup>a</sup> | 0.007 day /0.003 night (amol cell<sup>-1</sup> h<sup>-1</sup>)<sup>a</sup> or<br>0.9 day /0.4 night (pmol l<sup>-1</sup> h<sup>-1</sup>)<sup>a*</sup><br>0.0048<br>(amol cell<sup>-1</sup> h<sup>-1</sup>)<sup>d</sup><br>0.0026<br>(amol cell<sup>-1</sup> h<sup>-1</sup>)<sup>e</sup> | Undetectable<sup>a</sup><br>0.006<br>(amol cell<sup>-1</sup> h<sup>-1</sup>)<sup>e</sup> |
| **Glucose**<br>(<sup>14</sup>C-glucose) | 45.7<br>(pmol l<sup>-1</sup> h<sup>-1</sup>)<sup>a</sup> | Undetectable<sup>a</sup> | Undetectable<sup>a</sup> |

a This study
b Determined in the subtropical North Atlantic Ocean. Duhamel et al., (3).
c Determined in the subtropical and tropical Northeast Atlantic Ocean. Jardillier et al., (4).
d Calculated on the basis of the reported data. Determined in the South Pacific Ocean. Duhamel et al., (11).
e Determined in the Atlantic Ocean. Muñoz-Marín et al., (14)
\*Using cell abundances

*Prochlorococcus* represented 2.9 ± 1.3% of the glucose assimilation by the whole community (n = 20), with 3.4 ± 1.4% during daylight (n = 12) and 2.1 ± 0.8% during the night (n = 8) (Table 1). On average, the assimilation of carbon from glucose by the whole microbial community represented 0.63 ± 0.2% of the carbon fixed by primary production. The assimilation of carbon from glucose by *Prochlorococcus* represented circa 0.05 ± 0.02% of their carbon fixed by primary production (Table 1).

The ambient glucose concentration and the assimilation of glucose was initially measured using ^14^C-glucose as a potentially better tracker of the fate of carbon from glucose amendments. The ambient glucose concentration was 0.9 ± 0.1 nmol l^−1^ (n=5) and 0.5± 0.2 nmol l^−1^ respectively using ^14^C-glucose during the HOT 295 and 298 cruises. Taking both the ambient and added glucose concentrations into account, the ^14^C-glucose assimilation by the whole community showed the same pattern as the ^3^H-glucose assimilation, although varied by approximately a factor of 2.6 (27.7 to 73.9 pmol ^14^ C-Glc l^−1^ h^−1^), with on average 47.1 ± 12.6 pmol Glc l^−1^ h^−1^ (n = 20) (Table 1).

Due to the much lower specific activity of ^14^C-glucose compared to ^3^H-glucose, and the necessity to keep the glucose enrichments low at ~ 2 nM in these experiments, the radioactivity in *Prochlorococcus* sorted cells was not significantly different than in blank samples, and ^14^C-glucose assimilation cannot be reported for *Prochlorococcus*.

### Glucose effect on the bacterial community composition: 16 S rRNA sequences

We measured shifts in microbial community composition following glucose enrichment based on 16S rRNA gene tag sequencing. In all samples, the total and picoplankton communities were dominated by Alphaproteobacteria (mean 40% of total sequences in each sample vs. 3% for all other Proteobacteria combined), in particular the SAR11 clade, and cyanobacteria (38%), mainly *Prochlorococcus* (36%) (Fig. S6 and Supplementary Information). ASVs with unknown phylum were rare (<0.01%; Fig. S6). The remaining ASVs were mainly from Bacteroidetes (Flavobacteria), Actinobacteria (the OM-1 clade bacteria like *Candidatus* Actinomarina), Planctomycetes and Euryarchaeota (Fig. S6 and Supplementary Information).

There were no large shifts in the relative abundances of these taxa in response to glucose amendments over the 24 h incubation period based on NMDS and PERMANOVA analyses of Bray-Curtis distances between samples (Fig. S7; Supplementary Information). Thirty-three ASVs had significant (*p*<0.05) abundance changes in response to glucose but they were rare community members in each sample (< 0.1%; Supplementary Information, Fig. S8 and Table S3.

### High- and low-light Prochlorococcus photosystem I and C fixation genes were highly transcribed

The microarray targeted 1200 genes from the dominant *Prochlorococcus* strains at Station ALOHA (Table S4). On average, 41% (488 genes) were transcribed at detectable levels in each sample and 65% (775 genes) across all samples (Table S5). Thus, the population was transcriptionally active over the diverse set of metabolic pathways represented on the microarray (Table S4 and S5). Indeed, on average 64 ±33% (± standard deviation, n = xx) of genes were detected for each of the 34 strains represented on the microarray (Table S6). Within each sample most transcripts belonged to high-light (HL) adapted ecotypes from unknown clades (HOT208_60m_813O14, HOT208_60m_813G15), followed by clades II (AS9606, MIT0604) and I (MIT9515, MED4; Fig. S9). Low-light (LL) strains were also detected in every sample primarily from clades I (PAC1, NATL1/2) and II/III (SS120, MIT0602), and less often from IV (MIT9313, MIT0303; Table S6). Generally, we observed higher number of detected genes and transcript levels for HL strains than for LL (Table S6, Fig. S10). The fraction of the transcription levels of the LLIV strains (Fig. 2) are likely overestimates because only 9–10% of LLIV genes on the microarray were detected (Table S6).

**Figure 2.**
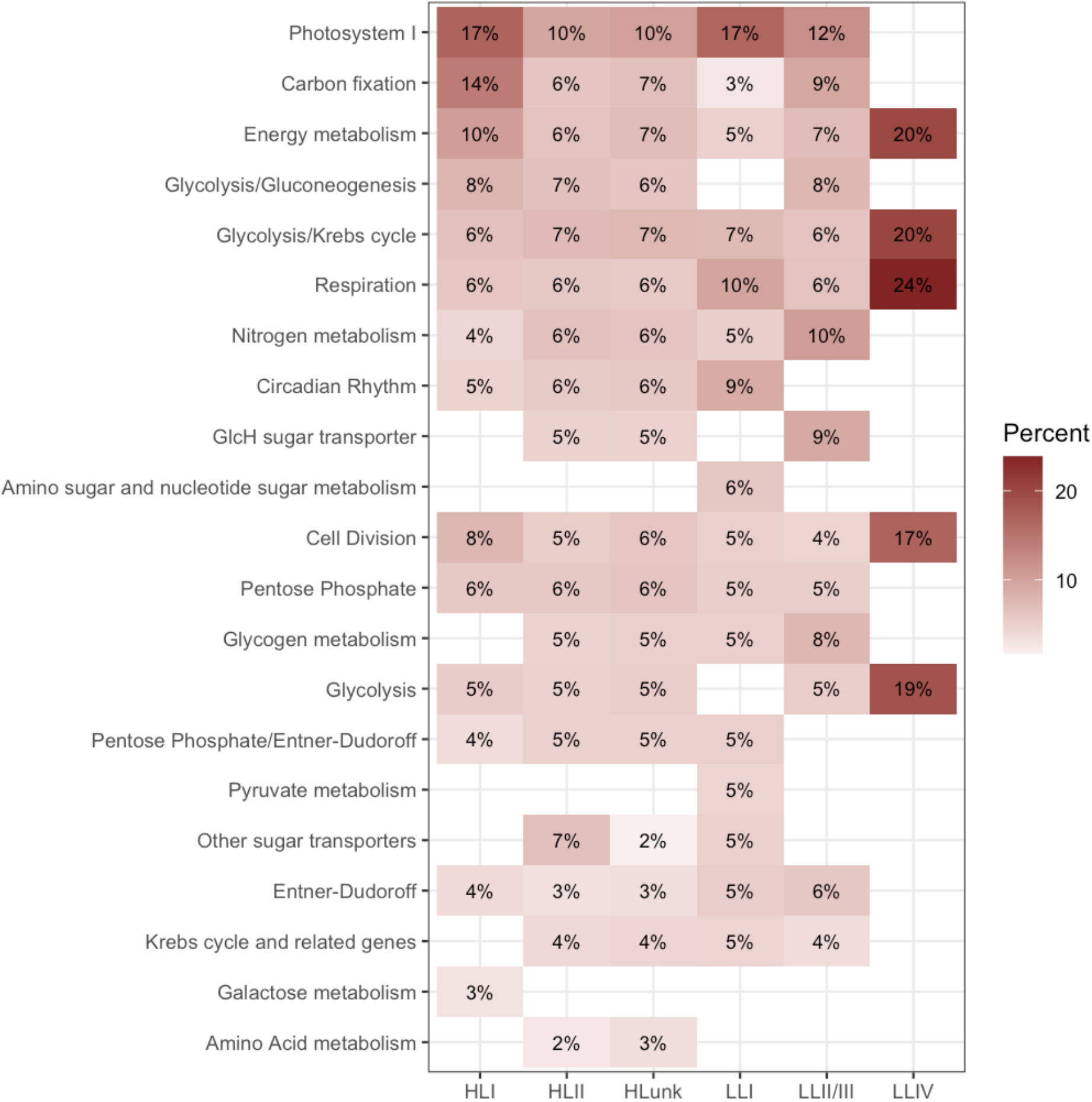
The heat map shows the transcription level across pathways (rows) in the controls for each *Prochlorococcus* clade (column). Within each control sample and for each clade, pathways were normalized for different gene counts by taking the mean transcript level for genes in the pathway. Pathway mean transcript levels were then summed and the percentages calculated. The percentages shown are the averages over control samples. Photosystem I had the highest percentages (excluding LLIV) and is therefore the top row. Empty cells indicate pathways that were not detected in the controls. The fraction of the transcription levels of the LLIV strains are likely overestimates because only 9–10% of LLIV genes on the microarray were detected.

Across all ecotypes, photosystem I (PSI) and C fixation pathways were highly transcribed in both control (Fig. 2) and glucose treatments (not shown), and notably transcription was higher for HL I compared to other HL clades. In contrast, LL clade I had higher levels of transcription for PSI and respiration but lower levels for C fixation when compared with other LL members (Fig. 2).

### Prochlorococcus expression patterns upon glucose addition

*Prochlorococcus* metatranscriptomes clustered primarily by day or night and secondarily by the hour of the day, which suggests that overall gene expression was more influenced by diel cycles than by glucose addition (Fig. S11). For most incubations, glucose treatments and matched controls had similar metatranscriptomes in the NMDS analysis. This suggests that if there were phase differences in diel expression between treatments and matched controls, despite mRNA having been fixed at approximately the same time, then the effects of the phase differences on metatranscriptomes were negligible. Moreover, the proportions of transcripts from each of the clades remained stable over the incubations from 4 to 24 h (Fig. S9). Therefore, any changes in the relative abundances of transcriptionally active HL and LL cells would also have had negligible effects on metatranscriptomes. Hence, we interpret differences in metatranscriptome positions in the NMDS between glucose treatments and matched controls as responses to glucose amendment. For the 4 h incubations that terminated in the night (20:00) or in day light (16:00), differences were small. We suspect that 4 h incubation was too short to observe a transcriptional response to glucose because larger metatranscriptome changes occurred for most of the incubations longer than 4 h. For example, metatranscriptome shifts were apparent in the NMDS for two of the three 12 h incubations that terminated at night (2D_12h_glucose and 12h_1glucose) and for the 24 h incubation that terminated in day light (2D_24h_glucose) in comparison to 4 h incubations that terminated at the same hour (16:00) (Fig. S11). However, some of the longer incubations did not show responses to glucose (for example 12h_2glucose and 24h_1glucose).

A total of 174 genes were significantly differentially expressed (DE) in response to glucose (Table S7). Expression changes for the 174 DE genes always exceeded 1.5-fold (mean 2.3-fold). Most DE genes (157 genes) were identified in the 12 h incubations that terminated in the dark at 4:00 (“2D_12h” for 2D_12h_glucose vs. 2D_12_control). Although we did not have replicates for 2D_12h, our DE tests borrowed information from all 15 microarrays to determine which genes had fold changes that were significant relative to their estimated gene variances (43, 45). HL *Prochlorococcus* had the majority of DE genes in 2D_12h (145 of 157 genes; Table S7), often transcript level increased within pathways involved in glucose metabolism (Fig. 3a and Table S7). For example, increases occurred for respiration (*coxAB*, *cyoC*), the pentose phosphate pathway (*tal, rbsK*), the Entner-Dudoroff pathway (*gdh*), glycolysis (*pgi*), glucose transport (*glcH*), and the Krebs Cycle (*fumC*). Increases also occurred for circadian rhythm genes (*kaiB*). Several carbon fixation genes increased (*rbcS*) but most decreased (*prk*), as did several genes involved in glycolysis/gluconeogenesis (*cbbA*, *pykF*) and energy metabolism (*atpABC*) (Fig. 3a and Table S7). A gene set enrichment analysis corroborated the HL increases for respiration, pentose phosphate pathway, and glucose transporter genes and the decreases for glycolysis genes (Supplemental Information). For LL, the 12 DE genes identified in 2D_12h included one *tal* (pentose phosphate) and one *coxA* (respiration) that decreased, whereas DE genes in these two pathways strictly increased for HL (Fig. 3b and Table S7).

**Figure 3.**
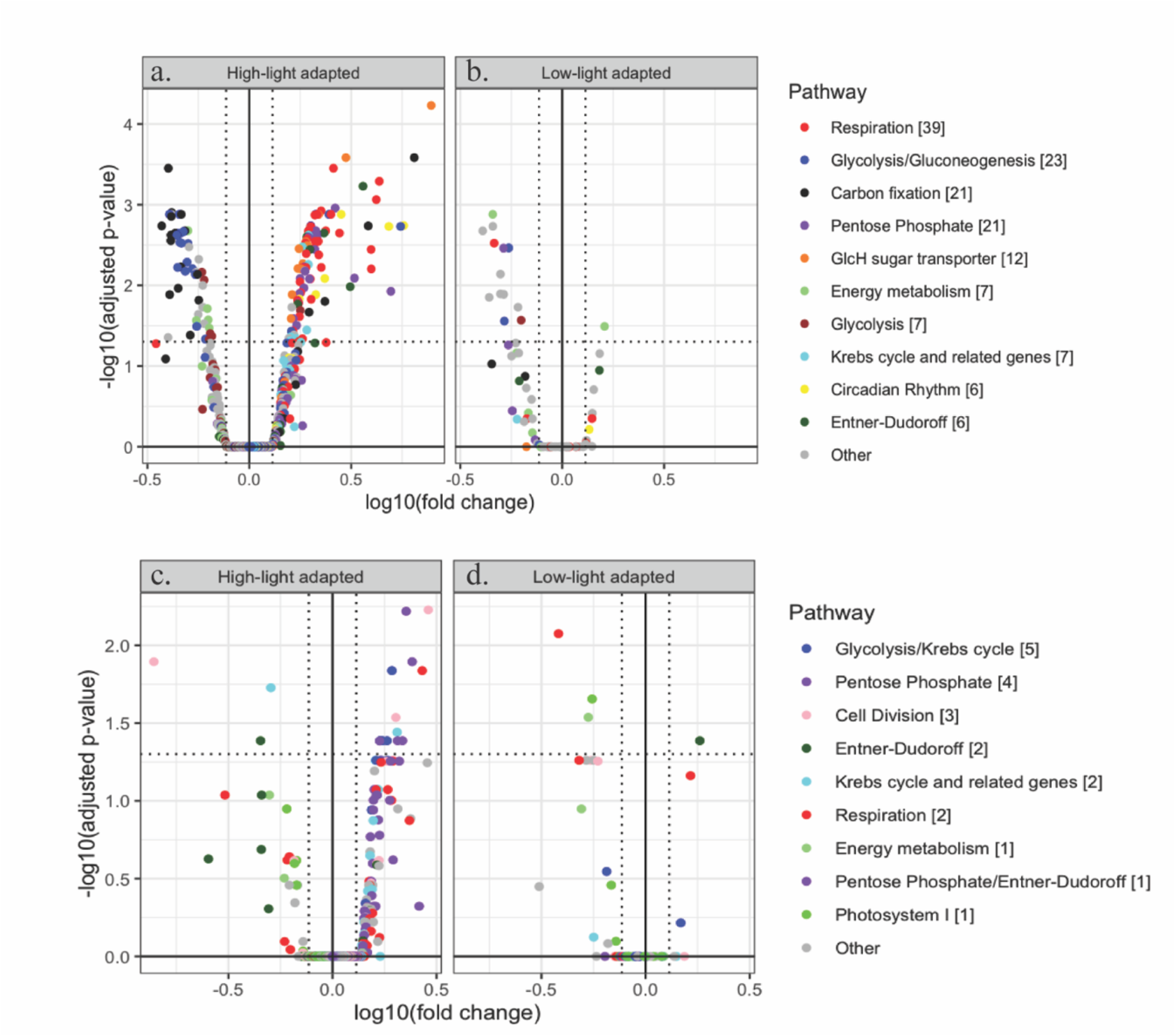
Top panel: A total of 157 DE genes from *Prochlorococcus* were identified in 2D_12h, the 12 h incubation that terminated at 4:00. Bottom panel: A total of 19 DE genes from *Prochlorococcus* were identified in 2D_24h, the 24 h incubation that terminated in day light at 12:00. The pathways and counts for the DE genes are in the legend. All detected genes from the pathways are shown in the plots separately for High-Light (a, c) and Low-Light (b, d) adapted *Prochlorococcus*. The DE genes are in the upper left and right sections of each plot, delimited by vertical dotted lines for fold changes > 1.3 and a horizontal dotted line for adjusted p-values < 0.05.

Sample metatranscriptomes clustered mainly by time of the day in both NMDS analysis, which used most detected genes, and in an analysis that used only the 157 DE genes in 2D_12h (Fig. S12). However, the 2D_12h samples did not cluster together, rather glucose addition resulted in transcript levels similar to those in the 2D_4h incubations (Fig. S12 [left]). This suggests that diel expression patterns were maintained for most incubations but were perturbed by glucose addition in 2D_12h. We visualized the response in a heat map (Fig. 4). For HL *Prochlorococcus*, the sample clusters were distinguished by pathways (mentioned earlier) that responded to glucose: elevated transcript levels for respiration and pentose phosphate pathway (gene cluster 1) and sugar transporters and Entner-Dudoroff (gene cluster 4), and decreased transcript levels for C fixation and glycolysis (gene clusters 2 and 3, respectively). Intriguingly, 5 *kaiB* genes were DE with increases from 2.1 to 5.7-fold, and all 5 of them (Circadian Rhythm genes in cluster 1) had their highest observed levels (in any samples) in 2D_12h_glucose. This might indicate that a change in diel regulation drove the transcription of 2D_12h_glucose to be more similar to the 2D_4h metatranscriptomes. LL *Prochlorococcus* had only 12 DE genes which mainly decreased in response to glucose amendment (in gene cluster 5, Fig. 4).

**Figure 4.**
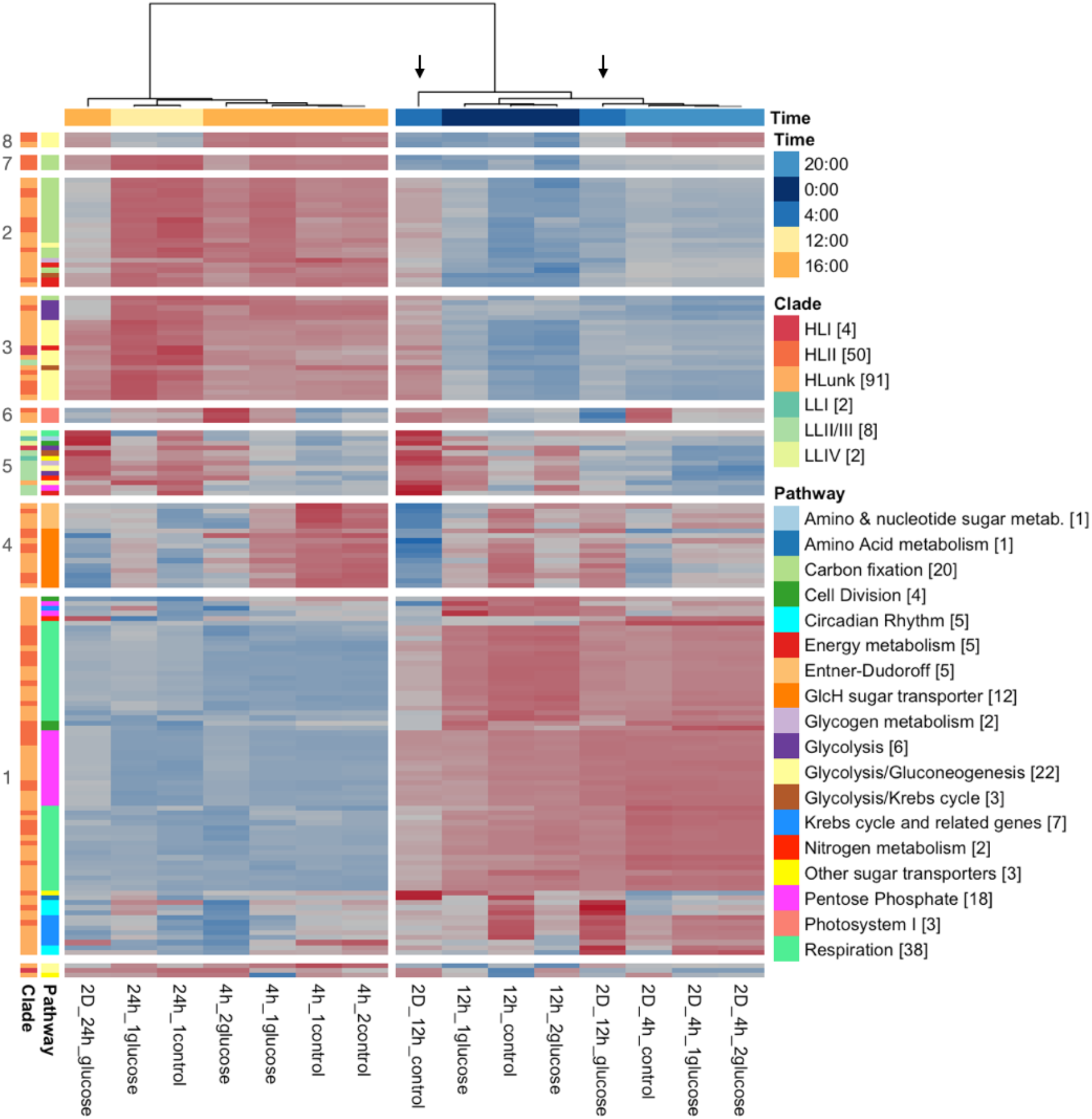
The heat map shows the 157 *Prochlorococcus* genes that were DE in 2D_12h glucose versus control (marked by arrows). Samples were hierarchically clustered (92) based on the Euclidean distances between 1 minus their Pearson correlations of the log_2_ transcript levels for the 157 DE genes. The two main sample clusters followed day and night and were significant (both had 97% support using multistep-multiscale bootstrap resampling with 10,000 bootstraps (93, 94). Genes (rows) had their log2 transcript levels standardized (mean=0, s.d.=1) prior to gene hierarchical clustering. Thus, transcription intensities (blue-red scale) should only be compared gene-wise, having lower transcription levels in blue or higher in red. Genes were hierarchically clustered using the same approach as the samples, and eight significant clusters were identified (>95% support). The bottom three genes are not members of the clusters. Row-side annotations include the numbers of DE genes from the category in brackets.

Most of the 19 DE genes identified in the 24 h incubations that terminated in the light at 12:00 (24h_glucose vs. 24h_control) were from pathways related to glucose metabolism (Fig. 3 c,d and Table S7). For HL, increases occurred for respiration (*coxC*), the pentose phosphate pathway (*opcA*), and pyruvate metabolism (*pdhB*), as well as for cell division (*minD*) (Fig. 3 c and Table S7). The respiration increases were corroborated by the gene set enrichment analysis (Supplemental Information). Few genes were DE for LL but they included *coxA*, which decreased, in contrast to strict increases for HL respiration genes (also in 2D_12h) (Fig. 3d and Table S7).

Only 1 DE gene was identified in the 4h incubations that terminated in the dark at 20:00 (2D_4h_glucose vs. 2D_4h_control), pyruvate dehydrogenase (*pdhA*) from the LLI strain NATL2, which decreased in the presence of glucose (Table S7).

## Discussion

### Prochlorococcus metabolism

*Prochlorococcus* abundances and cell size showed a clear diel cycle with increase in during the day and decrease during the latter half of the night as described in previous works (46, 47) (Fig. S5).

As expected, natural *Prochlorococcus* populations showed a pronounced diel cycle in carbon fixation, with undetectable values at night and 13.1 ± 8.5 nmol C l^−1^ h^−1^ (or 1.22 ± 0.66 fg C cell^−1^ h^−1^) fixed between noon and 16:00 (Fig. 1b, Table 2). Similar cell rates of carbon fixation by *Prochlorococcus* have been measured in the upper euphotic zone at Station ALOHA and in the Atlantic Ocean (3, 4, 8, 48–50) (Table 2).

Ambient glucose concentrations were in the nanomolar range, with averaged values of 1.1 ± 0.1 nmol l^−1^ (n = 4), similar to concentrations previously reported (11, 14, 20). Furthermore, the relative contribution of *Prochlorococcus* to total glucose assimilation was approximately 3.4 ± 1.4 % of the total glucose assimilation observed at Station ALOHA, very similar to previous reports from the North Atlantic Ocean (2.6–3.7%) (14) and the Western tropical South Pacific Ocean (~5%) (11). Based on the results obtained here, the glucose assimilation by *Prochlorococcus* represented a small fraction (< 1%) of total (inorganic + organic) C assimilation, similar to values previously reported (11, 14) (Table 1). It is worth noting that, in a previous study carried out in the Atlantic Ocean (14), the percentage of total glucose assimilation assigned to *Prochlorococcus* was overestimated due to errors in the calculation. The corrected data for glucose assimilation comparing the total C was also lower than 1%. Nevertheless, this percent could be underestimated if glucose is being used for energy rather than for biosynthesis, which would push the total amount of assimilated C from glucose above that determined from new biomass. Furthermore, glucose is one of the diverse dissolved organic C molecules pool present in the ocean (51, 52) that *Prochlorococcus* might be able to use (14, 53), which could make this percent much higher if we take into account all potential organic substrates.

It should be noted that the high affinity GlcH glucose transporter identified in *Prochlorococcus* is multiphasic, showing different Ks constants depending on the glucose concentration. (14). Therefore, if higher glucose concentrations become available in the ocean for whatever reason (54, 55), the transporter could work at higher glucose assimilation rates, and this could lead to a significant contribution of organic carbon assimilation by *Prochlorococcus*.

The total C assimilation by *Prochlorococcus* was determined comparing different tracers (^3^H-glucose versus ^14^C-sodium bicarbonate) since results obtained with ^14^C-glucose samples subjected to cell sorting were below detection limits. This could lead to underestimation of the C fraction assimilated from glucose due to a possible loss of ^3^H in exchange reactions with H_2_O, or to the fact that assimilated ^3^H can create problems of self-absorption (56, 57).

An interesting aspect of our results was the fact that *Prochlorococcus* showed a clear diel pattern in glucose assimilation with maximum values during the day (approximately 3-fold change) (Fig. 1d, Table 2). However, a contrasting diel pattern was observed for the whole community with higher values from midnight to early morning and low values at sunset (approximately 2-fold change).

Previous studies, carried out in the Pacific and Atlantic Oceans, showed similar per cell rates in daylight incubations (11, 14). Light stimulates the cyanobacterial assimilation of amino acid (8, 11, 58–61), DMSP (62, 63) and ATP (11, 60, 64); this has also been observed for the assimilation of glucose in natural populations of *Prochlorococcus* (11), where it is an active process (13, 15). However, this is the first study showing that glucose assimilation in natural *Prochlorococcus* populations follows a diel pattern. The fact that *Prochlorococcus* glucose assimilation rates peak during the light period while rates in the whole community peak during the night-early morning, could provide *Prochlorococcus* some advantages over the rest of the community. One of the advantages is the coupling of the energy produced by photosynthesis to the glucose assimilation, since it is actively transported (15). Coupling the light availability with cellular processes would facilitate adaptation to daily environmental changes (65). *Prochlorococcus* could thus be using some of the sugars that are lost by other microorganisms death and sloppy feeding by zooplankton during the day or other mortality (coevolved mutualism (66)). The fact that *Prochlorococcus* showed a different timing of glucose assimilation compared to the total population may also offer considerable fitness advantages over the competitors in “temporal niches” (67).

A similar difference in assimilation timing seems to exist for amino acids assimilation: in *Prochlorococcus* populations of the Atlantic Ocean, the maximum happens during the dark period (68); however, in heterotrophic bacteria studied in the Mediterranean sea, maximum leucine assimilation occurs around noon (69). In this regard, it would be worth investigating whether the assimilation of other organic compounds, such as ATP or DMSP, are also subjected to differential circadian rhythms in marine picocyanobacteria vs the total microbial community, which could be relevant in ecological terms.

Interestingly, previous studies on the diel rhythmicity of amino acid assimilation by *Prochlorococcus* in surface areas of the Atlantic Ocean showed maximal assimilation values at the beginning of the dark period, and minimal values around midday (68); this is almost exactly the opposite rhythm that we found for glucose assimilation in the same organism. This contrast is striking, especially if we consider that both amino acid and glucose assimilations are active processes, stimulated by light. A possible explanation for the difference might be based on the fact that amino acids are an important source for N in oligotrophic environments; since N is an essential element for the production of many cell compounds required before division, a maximum of amino acid assimilation at the beginning of the dark period might boost protein synthesis prior to *Prochlorococcus* cell division, as proposed by Mary and coworkers (68). By contrast, glucose can be directly used for general metabolic needs in *Prochlorococcus* (13), and therefore it would be more efficient to take up most glucose at midday, coupling the energy consumed by this process to the light photosynthetic reactions. Regardless of the difference of rhythms between glucose and amino acid assimilation in *Prochlorococcus*, the results show that not all light-stimulated assimilation processes are regulated the same way in marine picocyanobacteria.

Cell specific glucose assimilation by *Synechococcus* was previously determined in the Western tropical South Pacific Ocean and was similar to that of *Prochlorococcus* (0.006 amol Glc cell^−1^ h^−1^), likely attributable to *Synechococcus* larger biovolume (11) (Table 2). In the present study, the *Synechoccocus* population was also sorted after ^3^H-Glucose incubation but due to the low *Synechococcus* abundances at Station ALOHA (approximately 100-fold lower cell concentration than *Prochlorococcus*), results were not significantly different than the blanks. Still, it is possible that *Prochlorococcus* and *Synechococcus* compete for glucose. Experiments performed in laboratory cultures revealed that glucose transport in *Prochlorococcus* and *Synechococcus* displays multiphasic kinetic with high efficiency (calculated by dividing the assimilation rate by the K_s_ constant, between 0.01–20 μmol l^−1^) (15). A comparison of the assimilation efficiency demonstrated *Prochlorococcus* to be 7 times more efficient than *Synechococcus* (15), which could be an advantage for *Prochlorococcus* in oligotrophic areas where they coexist with *Synechococcus*, such as Station ALOHA.

### Effects of glucose enrichment

As anticipated from previous studies, the SAR11 clade (Proteobacteria) and *Prochlorococcus* were highly abundant in all samples at station ALOHA, followed by Bacteroidetes and Actinobacteria (70, 71).

Our results did not show differences in community composition after glucose enrichment (Fig. S6 and S7). It is possible that the incubation times were too short to see changes in the microbial community, or that the glucose concentration was too low to induce changes in the studied time. Higher abundance of *Prochlorococcus* upon addition of glucose and mannitol was observed in oligotrophic areas of the South Pacific (53); however, the authors used a 4,000-fold higher glucose concentration and longer incubation times than we did (0.4 mM and 78 h maximum vs 0.1 μM and 24 h maximum used in our study).

The population was transcriptionally active over the diverse metabolic pathways of the 34 strains identified. Many of the same *Prochlorococcus* strains detected in our results, were detected in previous studies also using Agilent microarrays at Station ALOHA (31). Furthermore, in both studies, photosynthesis and carbon fixation genes have been the most highly transcribed across all taxa and samples (this study and (31)).

Generally, we observed higher percentages of detected genes and higher transcript levels for HL strains than for LL (Table S6 and Fig. S10). Previous study at Station ALOHA also found much smaller proportions of LL clade transcripts relative to HL clades of *Prochlorococcus* at the surface (72). Our results might suggest that either HL strains had higher relative cell abundances (since samples were collected from the surface in our work) or were transcriptionally more active (73, 74), or both. However, if HL strains were transcriptionally more active than LL strains, we would expect even greater differences between the transcript abundances in each of the clades, since we are sampling at the surface where HL is more abundant (75–78).

Moreover, we found differences in the transcripts across the clades of HL and LL ecotypes, with high transcription level in PSI and C fixation pathways in HLI and LLI clades. As discussed above, these values might be related to the cell abundances of these clades, in fact a relatively high contribution of *Prochlorococcus* HLI and LLI in this North Pacific region has been observed previously in surface waters (77, 79–81). LLI strains are usually restricted to deeper depths at Station ALOHA when the water column is stratified, however contrary to other clades, LLI strains are present in the euphotic zone in mixed water (77).

*Prochlorococcus* strains showed the majority of the transcriptional changes after 12 h and 24 h after glucose enrichment (Fig. 3). Moreover, only one gene responded significantly in a 4h incubation (*pdhA* from NATL2 in experiment 2), which suggests that in most cases *Prochlorococcus* might require between 12 and 24 h from the moment that glucose is taken up until the transcriptional response for the glucose metabolism is detectable. Moreover, the surface light flux in the second experiment averaged 42.1 E m^−2^ d^−1^ versus 33 E m^−2^ d^−1^ during the first experiment (Table S1), which could explain most changes observed during the second experiment, where higher light could have stimulated the glucose assimilation.

A total of 173 genes were significantly DE in response to glucose after 12 h or 24 h incubations (Fig. 3 and Table S7). The effect of glucose enrichment on the transcriptome of HL strains showed increases for respiration (*coxAB*, *cyoC*), the pentose phosphate pathway (*tal, rbsK*), the Entner-Dudoroff pathway (*gdh*), glycolysis (*pgi*), glucose transport (*glcH*), the Krebs Cycle (*fumC*), pyruvate metabolism (*pdhB*) and cell division (*minD*). The largest transcriptional changes occurred after 12 h incubation in the genes encoding the glucose transporter (*glcH*) with almost 8-fold increase, small RUBISCO subunit (*rbcS)* with 6.4-fold increase and the glucose 6-phosphate isomerase (*pgi*) involved in glycolysis with 5.5-fold increase (Table S7 and Fig. 3). Furthermore, a gene set enrichment analysis corroborated the HL increases for respiration, pentose phosphate pathway, and glucose transporter genes and the decreases for other glycolysis genes (Supplemental Information).

It has been proposed that *Prochlorococcus* might use two pathways to metabolize glucose, the Entner-Dudoroff and the pentose phosphate pathways (12, 13, 15, 82) and small changes in gene expression and quantitative proteomics have been demonstrated upon glucose addition (13, 15). Our results show, for the first time in natural samples, that *Prochlorococcus* could utilize glucose by both pathways. Changes in the expression of one glycolytic enzyme (phosphoglucose isomerase, *pgi*) was also observed, as previously reported (15). Glycolysis is not active since *Prochlorococcus* lacks phosphofructokinase (82). However, even if it lacks the enzymes involved in the initial steps of glycolysis, this cyanobacterium still has a few genes which could be involved in glucose assimilation with the production of reducing equivalents and/or the production of ATP as a result of the metabolization of glyceraldehyde 3-phosphate and phosphoenolpyruvate (13, 82).

Periodicities of the transcripts of genes involved in physiological processes such as carbon fixation, energy metabolism, photosynthesis, respiration, pentose phosphate, cell division, and amino acid metabolism tracked the timing of its activities relative to the light-dark cycle, as previously described (83). We observed high *glcH* transcription levels during the night at 20:00 but mostly at 4:00 with the maximum transcript level (~8-fold change) (Supplemental Information and Table S7). The highest transcriptional changes of *glcH* during the night could indicate the synthesis of the glucose transporter in order to be ready during the day, when light is stimulating the assimilation according to the diel pattern in glucose assimilation.

An interesting result for HL strains was that glucose addition led to transcript level increases for the circadian gene *kaiB* (2.1-5.7-fold) 12 hours later at 04:00. A hypothesis is that glucose addition affects diel expression patterns. Zinser and coworkers previously described diel cycles for the HL strain MED4 in culture (83). A qualitative comparison to that work indicated that some of the changes we observed were similar but found earlier in the diel cycle than those described by Zinser. Diel shifts were supported by our DE and gene set enrichment analyses (usually both) for multiple pathways. The transcript levels of Photosystem I and energy metabolism (ATP synthases) genes decreased, and respiration (*coxAB*) and the pentose phosphate pathway (*tal*) genes increased, consistent with MED4 diel expression changes from midnight to sunrise. Although we did not observe *rbcL* decreases as would be expected with a shift to earlier in the cycle, we saw clear decreases from phosphoribulokinase (*prk*), another enzyme in the Calvin-Benson Cycle. Curiously, we would have expected *kaiB* to decrease but observed the opposite. However, a close look at Zinser et al. shows some variability in *kaiB* levels as they increased to a sunrise peak. Therefore, in comparing the work of Zinser and coworkers on diel patterns for *Prochlorococcus* sp. MED4 with our work, we speculate that glucose addition might have delayed the diel expression patterns in the natural *Prochlorococcus* population studied.

Several studies have shown that circadian clocks are connected to cyanobacterial metabolism (84–86). Moreover, it has been found that in the presence of sugars the circadian clock acts as a dynamic homeostat responding to the carbohydrate signals (87) and, in other cyanobacteria, blocking the clock-resetting effect of a dark pulse (85). Our results suggest that glucose assimilation affects the circadian transcriptional machinery in *Prochlorococcus*, in good agreement with the studies cited above. The sudden availability of glucose could be an important event for the metabolism of *Prochlorococcus*, especially under conditions of darkness or very low light (88–90), justifying a change in the response to light rhythms.

Finally, we observed differences in the transcription profile between the different ecotypes. Few genes were DE in LL *Prochlorococcus* strains, but those that were showed different patterns after glucose addition compared to HL strains. In fact, the LL ecotype was clustered in the heat map as an independent gene group (gene cluster 5). Genes from LL clades involved in pentose phosphate and respiration decreased after glucose addition, whereas genes in the same pathways increased in HL strains after 12 and 24 h of glucose addition (Fig. 3b,d). Other studies have also found that coexisting *Prochlorococcus* populations respond differently upon nutrients addition (91). Subpopulations with different abilities to utilize glucose even at the same depth could be explained by many factors like competition, environmental conditions or genetics. In fact, there is a diversity of kinetics in glucose assimilation in the *Prochlorococcus* strains (15), which could suggest that the glucose assimilation has been subjected to diversification along the *Prochlorococcus* evolution.

Overall, our results indicated that *Prochlorococcus* shows synchronous timing in gene expression and in glucose assimilation presumably coupled to the light cycle. Diurnal glucose assimilation allows *Prochlorococcus* to optimize glucose assimilation by using ATP made during the light period, coupling this process to photosynthesis. Furthermore, it also could provide some advantages over the rest of the community, which showed a different timing for glucose assimilation. This hypothesis might be related to the possible glucose-induced delay in the transcriptional rhythms suggested by some of our results.

The relative contribution of the different metabolic pathways to metabolize glucose in different subpopulations of *Prochlorococcus* should be further investigated to understand the impact of mixotrophy on the marine cyanobacterial populations and their consequences for global biogeochemical cycles.

## Declarations

## Acknowledgments and funding

We are grateful to the captain and crew of the R/V Kilo Moana for their essential support.

M.C.M.-M. was supported by a Marie Curie International Outgoing Fellowship within the 7th European Community Framework Programme (FP7-PIOF-GA-2013-625188). S.D. was funded by the National Science Foundation (OCE-1434916). K.B. and D.M.K. were funded by support of the Simons Foundation (SCOPE award 721252) and the National Science Foundation (OCE-1756517; A.E. White, PI). M.C. M.-M, J.D. and J.M. G.-F. were funded by the Spanish Ministry of Economy and Competitiveness (BFU 2016-76227-P).

## Availability of data and material

Microarray data have been deposited at NCBI Gene Expression Omnibus (GEO) under accession number GSE154594. The 16S raw sequences have been deposited at Sequence Read Archive (SRA) with the BioProject ID PRJNA758505.

## Authors’ contributions

M.C.M.-M. and S.D. designed the study. M.C.M-M. sampled, designed and performed the molecular approaches (transcriptomic and metagenomic analysis) and J.M analyzed the 16 S and transcriptomic data and performed the corresponding figures. S.D. and K.B. sampled, designed and performed the radioassays (cell-specific and bulk, respectively) and analyzed the data. M.M.M., S.D., K.B., J.M., J.D., D.K., and J.M.G.-F. drafted and edited the manuscript and figures. All authors read and approved the final manuscript.

