## Supplemental Information for "Differential timing for glucose assimilation in *Prochlorococcus* and coexistent microbial populations at the North Pacific Subtropical Gyre"

**This file includes:**

Supplementary Information
Figs. S1 to S12
Tables. S1, S8 and S9
Captions for Tables S1, S2, S3, S4, S5, S6, S7, S8, S9 and S10.
References for supplementary information

**Additional data (separate files)**

Tables S2, S3, S4, S5, S6, S7 and S10 (excel files)

### Supplemental Information

Extended description of Material and Methods described in the main text.

*16S rRNA analysis*

A total of 718 ASVs were identified within the 16S rRNA data sets (Methods, Table S2). The total numbers of reads assigned to ASVs were similar across samples (mean 132,955 reads in ASVs, range 101,003 to 156,099). Rarefaction curves (with vegan R package function rarefaction) showed that ASV counts saturated well before 101K reads for all samples. Therefore, sequencing depth in each sample was sufficient to capture the richness of the microbial community. Rarefying to 101K reads, samples had a mean of 404 ASVs (s.d.=33).

Proteobacteria and cyanobacteria dominated all samples and had stable relative abundances (means were 0.43±0.04 and 0.38±0.02, respectively) despite different sample collection dates, treatments, and incubation times (Fig. S6). Alpha-proteobacteria dominated and Delta- and Gamma-proteobacteria were rare (mean relative abundances were 0.40, 0.02, and 0.01, respectively). *Prochlorococcus* dominated cyanobacteria (mean 0.36) and *Synechococcus*, the only other identified cyanobacterial genus, was rare (<0.01).

NMDS analysis of the 16S rRNA ASV abundances was performed using vegan (Fig. S7). First, the ASV count matrix was scaled to normalize sequencing depths (decostand ('total')). Then metaMDS was used with Bray-Curtis dissimilarities and auto-transformation disabled, but default parameters otherwise. The NMDS stress was 0.08. Samples did not cluster by treatment or incubation time (Fig. S7.Top panel). However, samples clustered weakly by experiment (experiment 1 samples 7, 3, 19, and 23 in Fig. S7. Bottom panel) indicating small differences between communities collected on October 6 at noon (experiment 1) versus on October 7 at 16:00 (experiment 2). Indeed, the relative abundance of *Bacteroidetes* was higher in samples collected on October 6 (0.12) than on October 7 (0.08; $p<0.01$, Welch two-sample $t$-test).

To identify ASVs that had statistically significant abundance changes in response to glucose, we used the edgeR and limma packages (1, 2). EdgeR transforms sequence count data so that they can be used with the same limma functions as in the microarray differential expression (DE) analysis. Counts for the 718 ASVs were imported into an DGEList, and then rare ASVs were removed as described in (3) and implemented in filterByExpr, leaving 678 ASVs. Normalization factors to correct for different sequencing depths were calculated using the TMM approach described by (4) and implemented in calcNormFactors. Sequencing depths were similar (101-156K reads) so the normalization factors had a narrow range (0.93 to 1.10). The normalized ASV count matrix was transformed to log counts per million (cpm). Next, we used the limma-trend method (5, 6) to identify differentially abundant ASVs, with steps nearly identical to those in the DE analysis. Linear models for the ASV abundances were created (lmFit) and empirical Bayesian inference performed with moderated $t$-statistics relative to a 1.2× fold change (treat with trend=TRUE). ASVs that had fold changes significantly larger than 1.2× ($p<0.05$) had to pass a final test: The ASV had to have >50 reads in at least one of the samples compared, for example, in at least one of the four samples in the 2D_12h_glucose vs. 2D_12h_control comparison. Although an ASV with 50 reads would comprise only 0.04% of the reads in a sample (mean; max 0.05%), that ASV would be more abundant than 41% of the other ASVs detected in the sample (on average).

A total of 33 differentially abundant (DA) ASVs were identified in five comparisons of matched glucose vs. controls (Table S3). None had more than a few hundred reads in any of the compared glucose and control replicates (maximum of 363 reads; Fig. S8). All of the DA ASVs had 0 reads in one of the conditions compared and >0 reads in the other condition (Table S3), which indicated that the ASV changed from undetected to detected. Although detected, the DA ASVs had small relative abundances (medians < 0.001 and maximum = 0.005; Fig. S8) In comparison, the $100^{th}$ most abundant ASV in each sample had relative abundance slightly >0.001. Moreover, PERMANOVA analyses (with vegan adonis2 using Bray-Curtis dissimilarities and 999 permutations, blocked by experiment) indicated that glucose addition did not have a significant impact on community composition ($p>0.05$, whether samples were represented by ASV or phylum abundance profiles). Therefore, we interpret the abundance changes in response to glucose as too small to be biologically meaningful. This is consistent with the NMDS results which did not show samples clustering by treatment.

*Differentially expressed pathways*

The gene differential expression analysis identified individual genes in specific *Prochlorococcus* strains that responded to glucose addition. We also used a complementary approach, an Ensemble of Gene Set Enrichment Analyses (EGSEA;(7)), to identify differentially expressed pathways from HL and LL *Prochlorococcus*. EGSEA identifies sets of genes that collectively show significant differential expression based on a consensus of 12 GSEA algorithms. As in a previous study by Shilova and colleagues (8), gene sets were defined by pathway and phylogroup, HL or LL in the present work (Table S8). Each pathway included genes that would change in the same direction and thus reinforce any signal detected by EGSEA. Differentially expressed pathways for HL and LL were tested in the same treatments versus controls as in the DE gene analysis (Table S7). The *p*-values from the 12 GSEA algorithms were combined using Wilkinson's method (9) and then corrected for multiple testing using the approach of Benjamini and Hochberg (10). Adjusted *p*-values < 0.01 were significant.

In comparison to the DE results in experiment 2 after 12 h for HL *Prochlorococcus* (main text), EGSEA corroborated increases for respiration, pentose phosphate pathway, and sugar transporter (*glcH*) genes, as well as the decreases for *pykF* genes in glycolysis (Table S9). DE results that were not corroborated by EGSEA were due to the adjusted *p*-value exceeding 0.01. However, the direction of change identified by EGSEA was always consistent with the reported DE genes. For example, Entner-Dudoroff had 5 *gdh* gene targets (from 5 distinct strains) with transcript level increases (1.7-3.6-fold; Table S7), whereas EGSEA found that collectively the 23 genes detected in the pathway (Table S8) did not change significantly (*p*=0.20) despite an average 1.6-fold increase. Similarly, for the Krebs Cycle the DE analysis identified 6 *fumC* and 1 *ppc* targets (1.6-2.0-fold increases), whereas EGSEA found that the 40 detected genes did not change significantly (*p*=0.02; average 1.4-fold increase). EGSEA also found insignificant increases for the 25 detected RuBisCO genes (*p*=0.02; average 1.5-fold increase), whereas the DE analysis found 4 *rbcS* genes that increased (2.0-6.5-fold). Interestingly, EGSEA identified small but significant decreases from photosystem I genes (average 1.3-fold decrease) even though only 3 of the 40 detected genes decreased (and none increased) in the DE analysis. These results underscore the robustness of the DE results by a complementary approach, EGSEA.

Note that the "Circadian rhythm" gene set included only *kaiC* genes (Table S8) and does not capture the *kaiB* increases described in the main text. The full EGSEA results are in Table S10.

**Titles and legends for the supplementary figures**

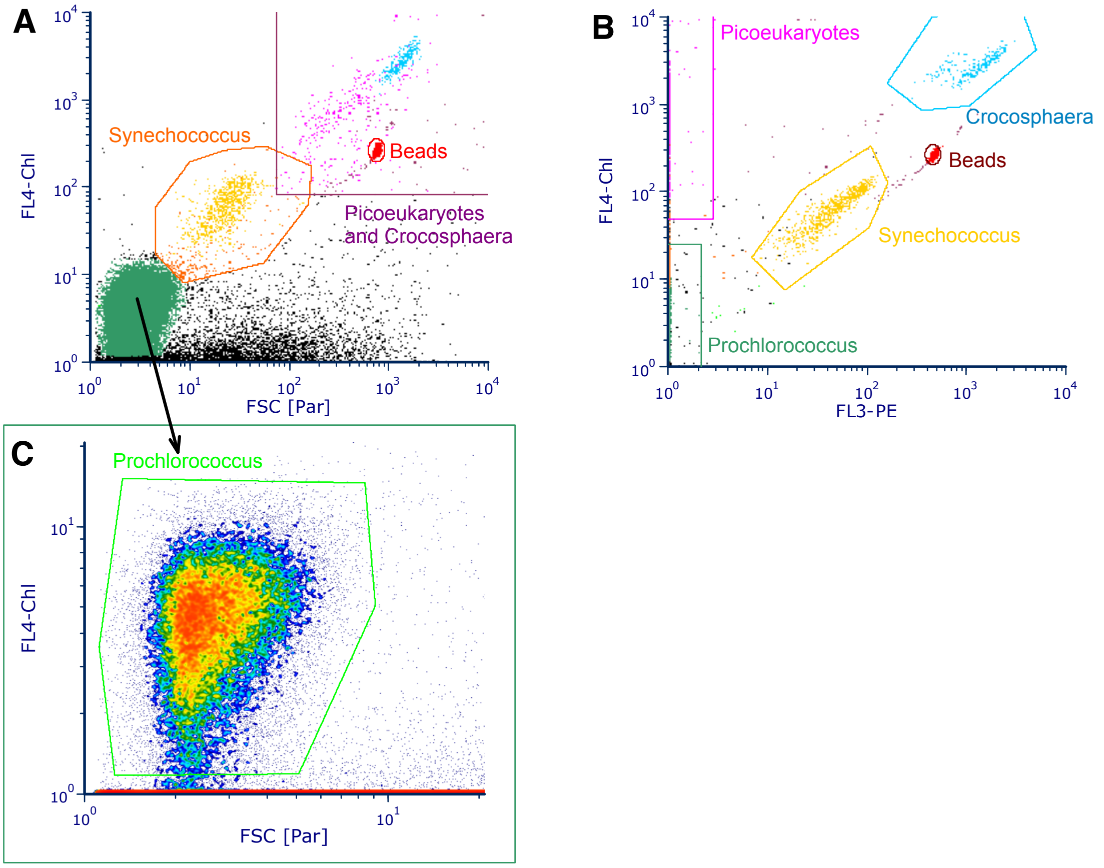


**Figure S1.** Examples of cytograms to illustrate the gating strategy employed to differentiate the picophytoplankton populations in our samples. A) Color dot plot of red fluorescence (FL4-Chl) vs forwards scatter (FSC[PAR]) showing the primary gates; B) Color dot plot of red fluorescence (FL4-Chl) vs orange fluorescence (FL3-PE) showing the secondary gates used to refine overlapping populations in A; C) Density plot showing the primary gate used to identify *Prochlorococcus* and differentiate this population from the instrument noise (note the different axis scales used to zoom on the population of interest).

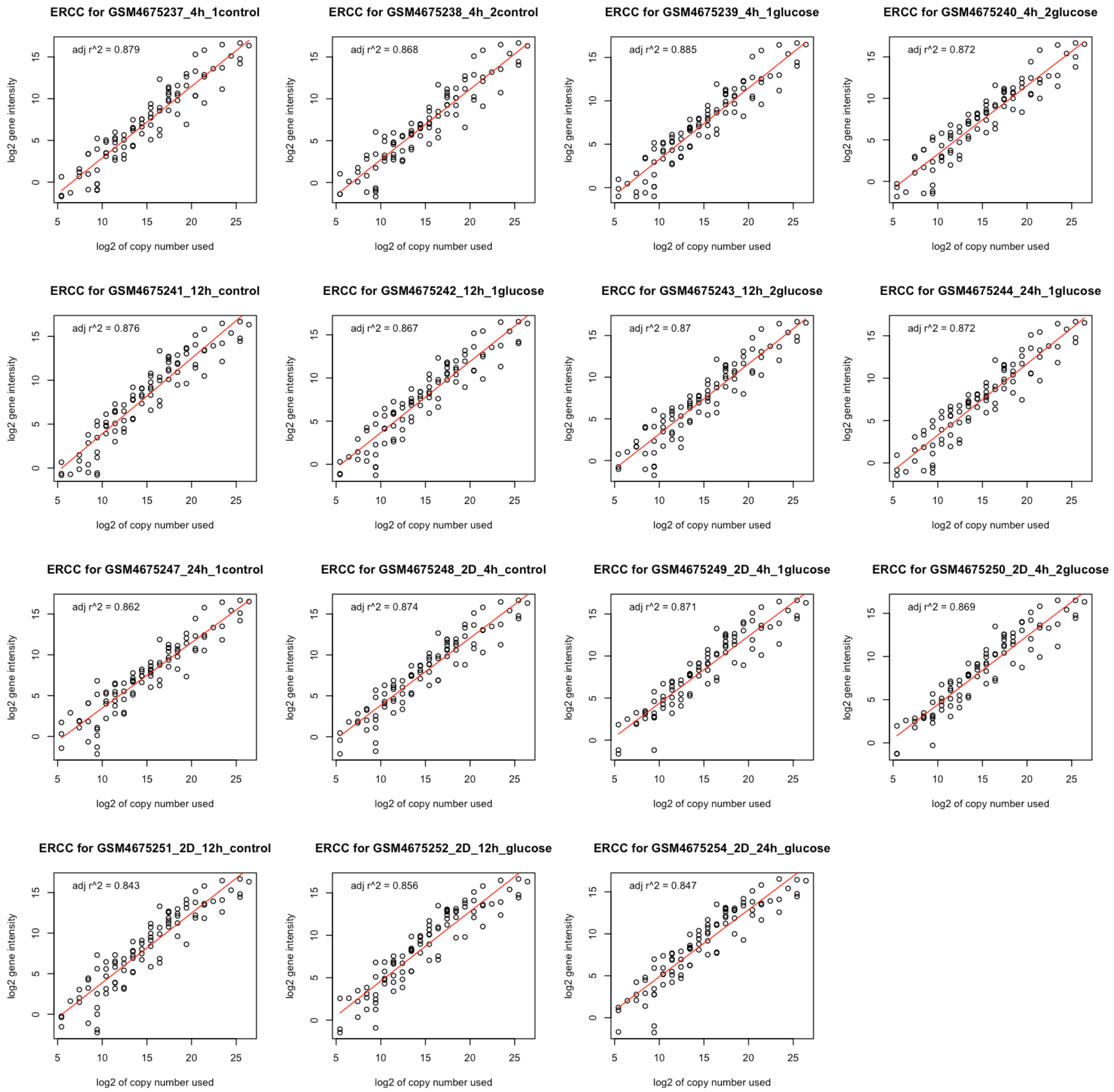


**Figure S2.** Hybridization signal for the ERCC mRNA spike-in mix (External RNA Control Consortium (Ambion®)) (a) added to eight samples (24hC-1.1, 2D_4hG-1.1, 2D_4hG-1.2, 2D_4hC, 2D_12hG, 2D_12hC, 2D_24hG, 2D_24hC) in the 8x60K array slide, (b) eight samples (12hG-1.1, 12hG-1.2, 12hC, 24hG, 4hC-1.1, 4hG-1.1, 4hC-1.2, 4hG-1.2) in the second 8x60K array slide.

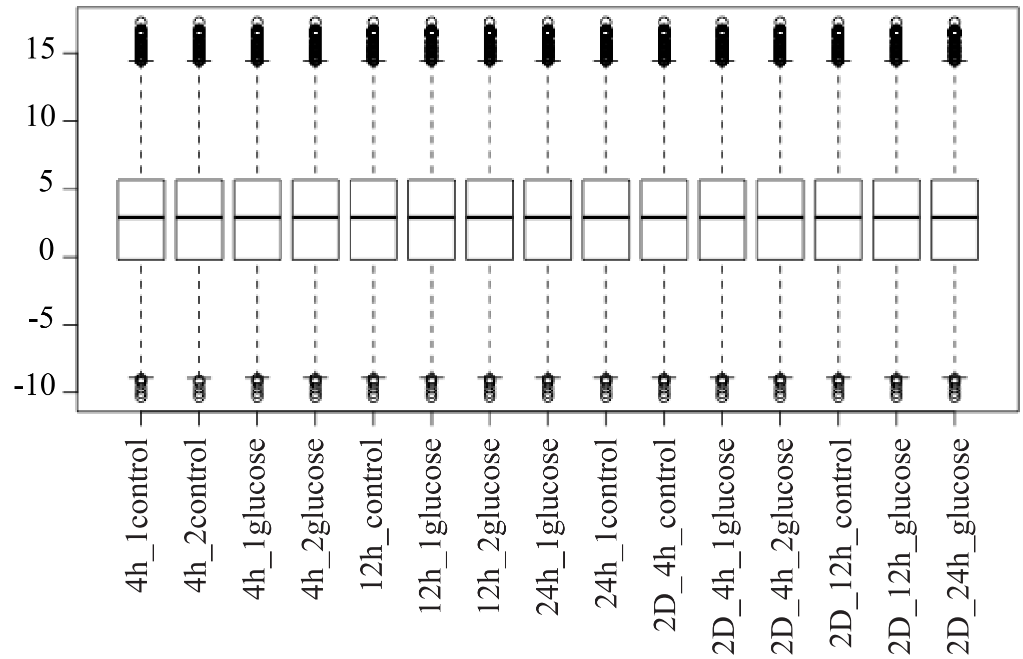


**Figure S3.** Boxplot of the hybridization signals ($log_2$) for all 15 *Prochlorococcus* microarrays after quantile-normalization of the ~14K experimental probes.

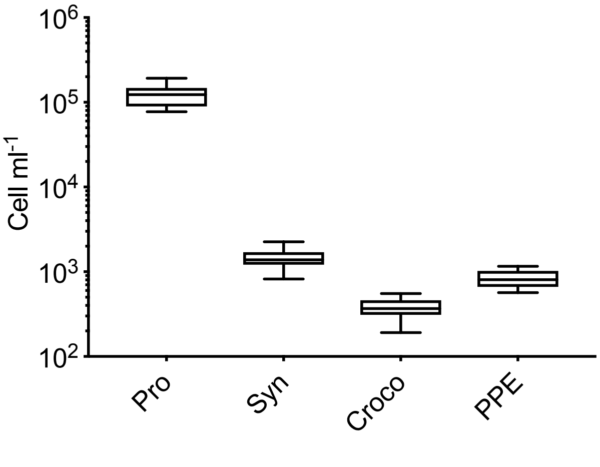


**Figure S4:** Boxes and whiskers plot of picophytoplankton cell abundances (cell $ml^{-1}$): *Prochlorococcus* (Pro), *Synechococcus* (Syn), *Crocosphaera* (Croco) and picophytoeukaryotes (PPE). The box extends from the 25th to 75th percentiles and the line within the box represents the median. Whiskers represent the minimum and maximum values.

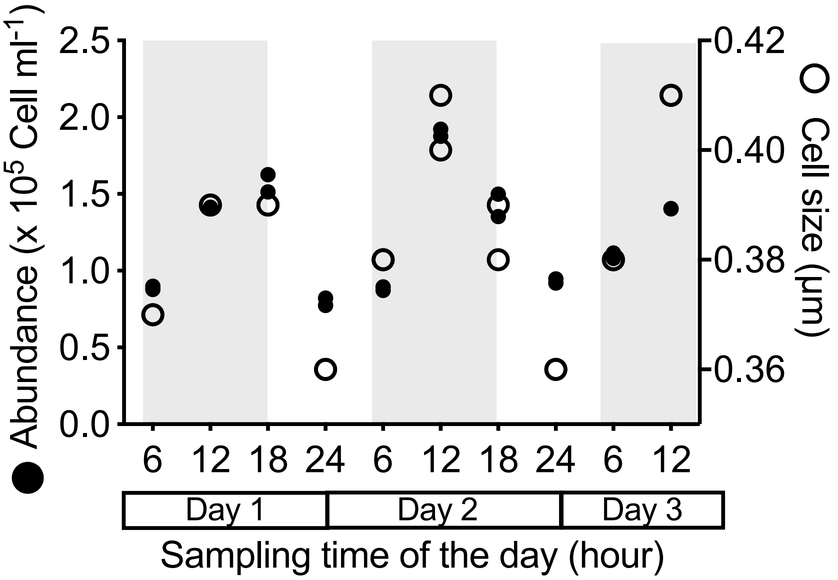


**Figure S5.** Cell abundances ($10^5$ cells $L^{-1}$, black dots) and cell size measurements (µm, white dots) during the three sampling days. The shaded area represents the dark period.

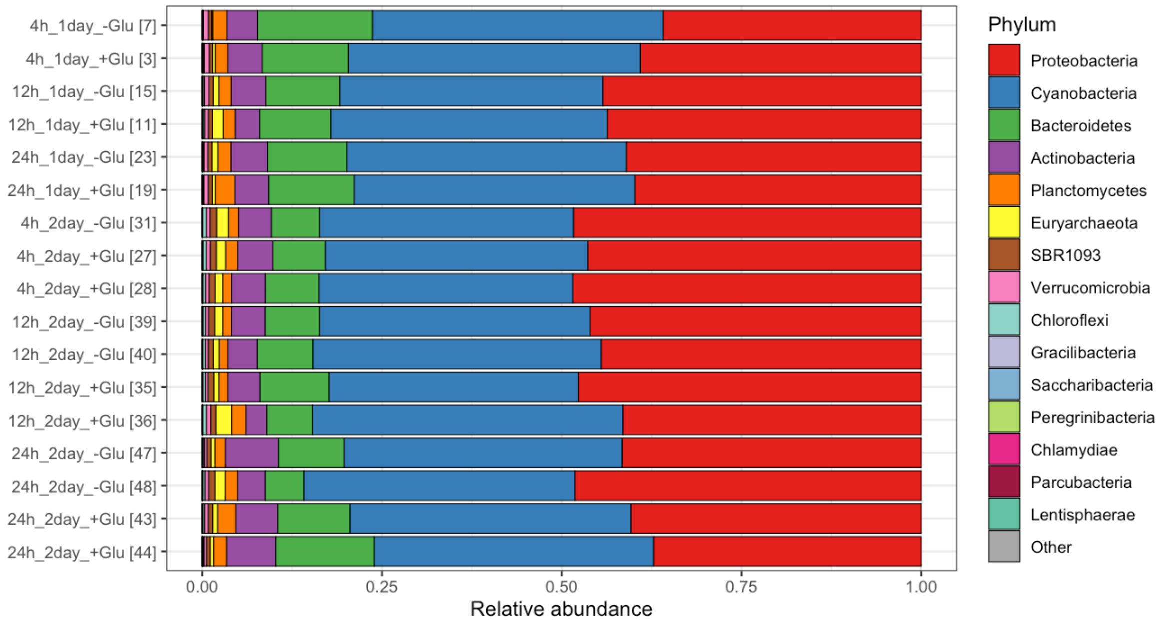


**Figure S6.** Phylum relative abundances in the 16S rRNA data sets. The 718 ASVs were binned by phylum. Rare ASVs or those with unknown phylum were binned in Other. Samples are grouped by experiment (first 6 are experiment 1), then incubation time, and then treatment. Row labels include sample ID numbers in brackets.

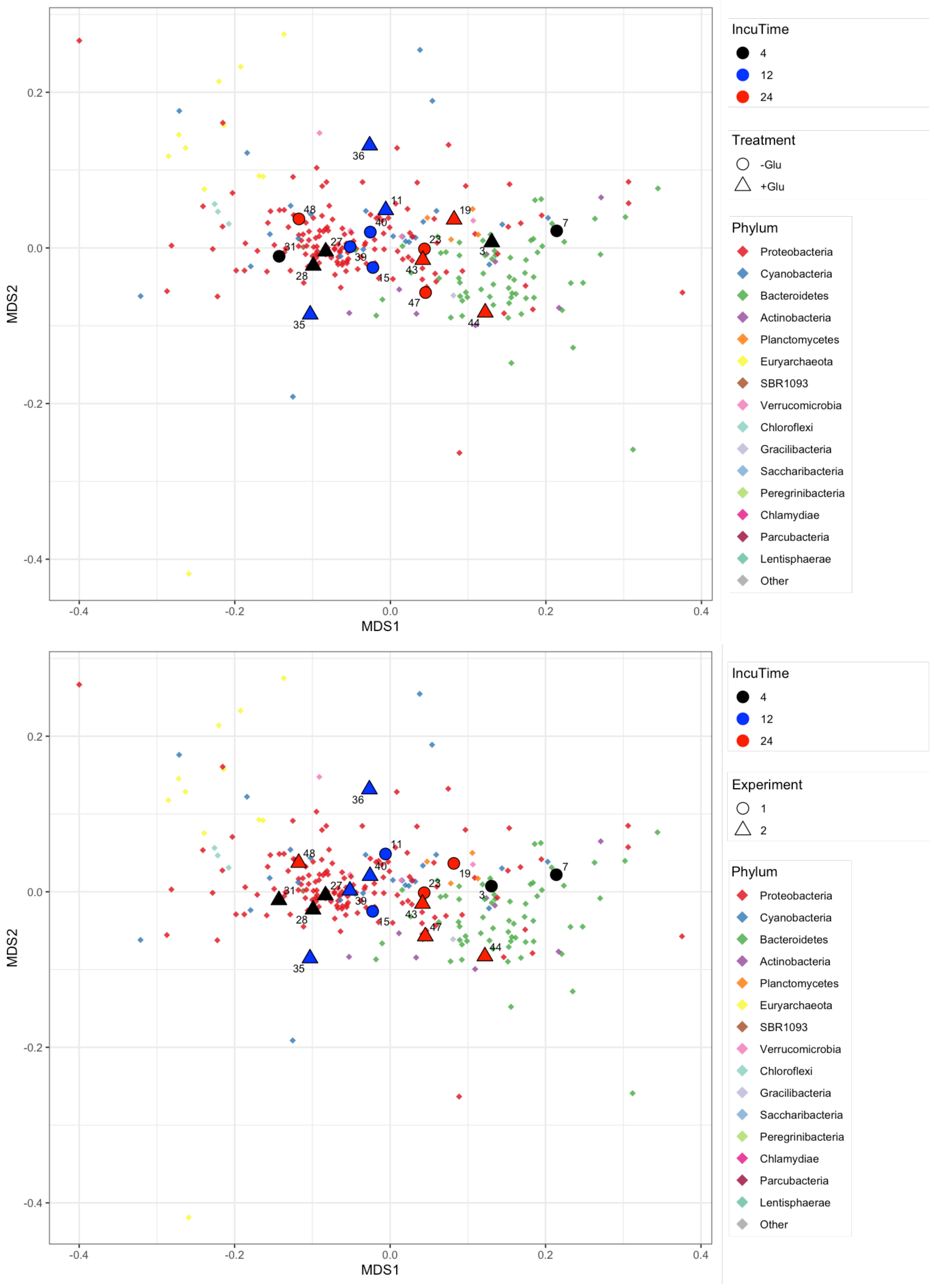


**Figure S7.** NMDS of the 16S rRNA samples represented by their ASV abundance profiles (stress 0.08). ASV coordinates are weighted (by `metaMDS`) by their abundances in the samples. ASV colors indicate phyla, which are ordered by total abundance in the legend. For legibility, only 270 non-rare ASVs are shown, defined as those with >5 reads in all samples, or >50 reads in at least 8 samples, or >100 reads in at least 3 samples. Top panel: Shapes indicate treatment (glucose and non-glucose addition) after 4h (black), 12h (blue) and 24 h (red) incubation. Bottom panel. Shapes indicate experiment 1 or 2, with samples collected at noon or 16:00 respectively, after 4h (black), 12h (blue) and 24 h (red) incubation.

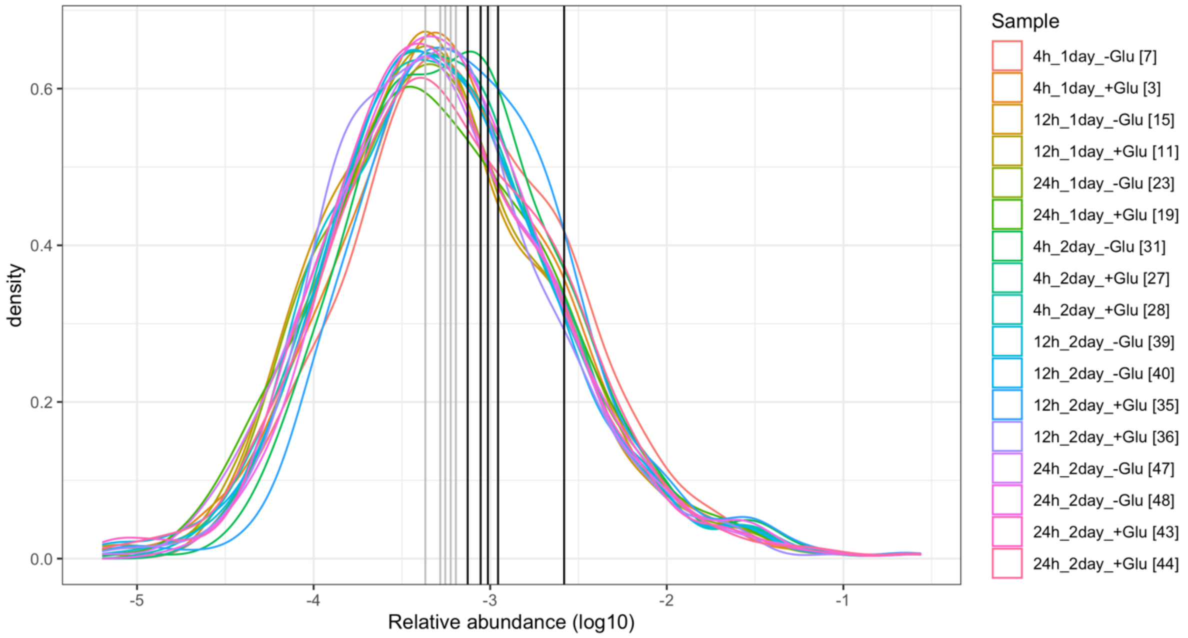


**Figure S8.** For each sample, the distribution of relative abundances for detected ASVs is shown. Vertical lines indicate the maxima (black) and median (gray) relative abundances of the DA ASVs identified in any of the five comparisons, with black for maxima and gray for medians.

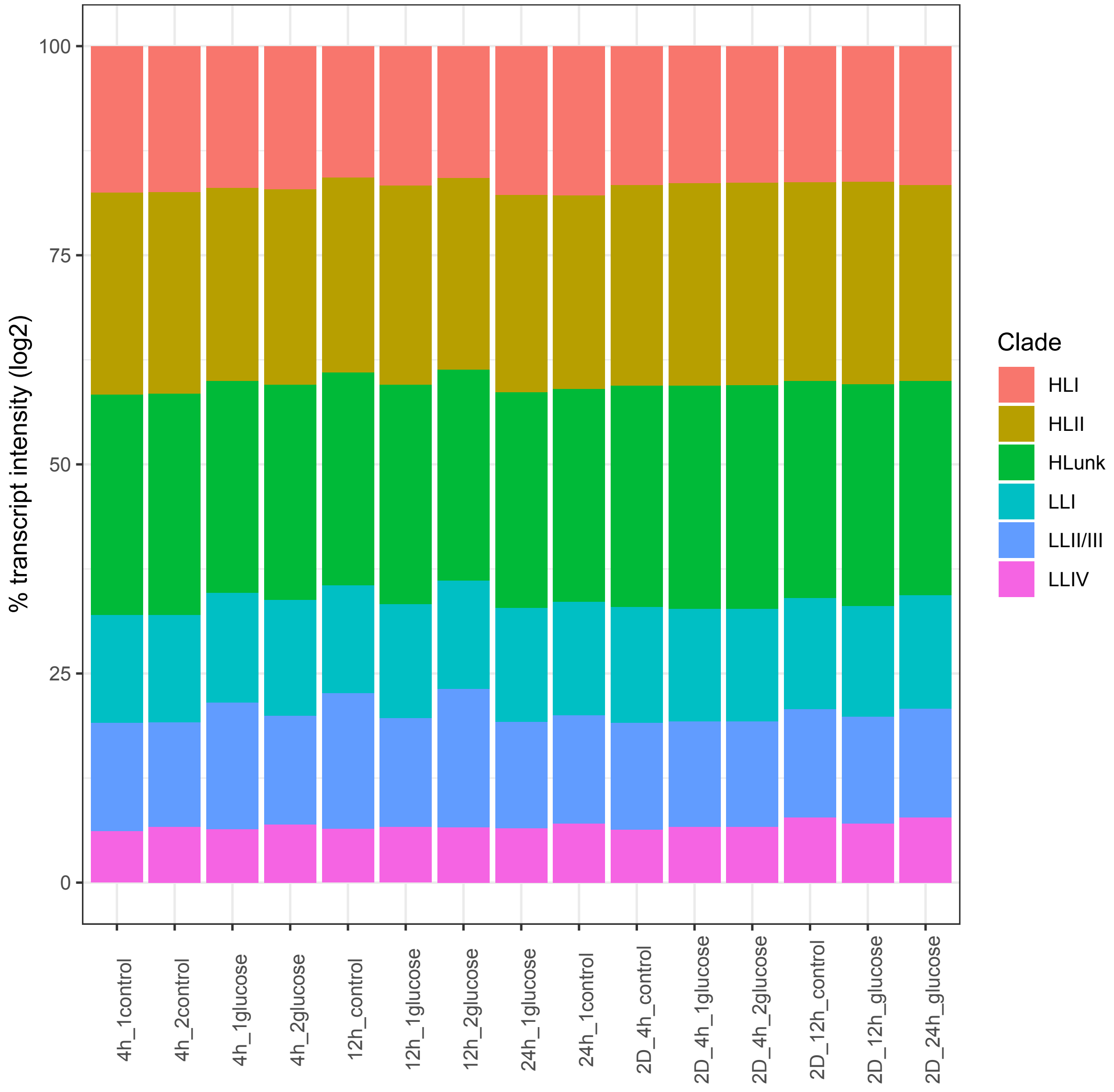


**Figure S9.** Total transcript levels (normalized, $log_2$) for each *Prochlorococcus* clade across the samples. Sample labels indicate the incubation time (4h, 12h and 24h) in the presence of glucose or in the control treatment, and the replicate (1 or 2). Samples from the second experiment ("2D") were biological replicates.

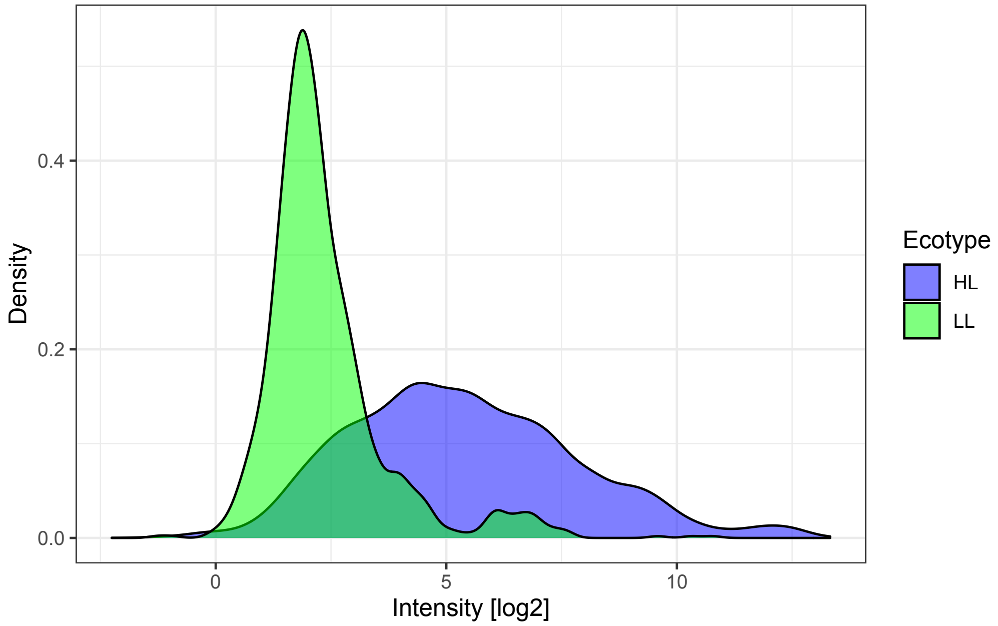


**Figure S10.** Distribution of normalized $log_2$ transcript levels for detected genes from High Light (HL) and Low Light (LL) ecotypes of *Prochlorococcus*.

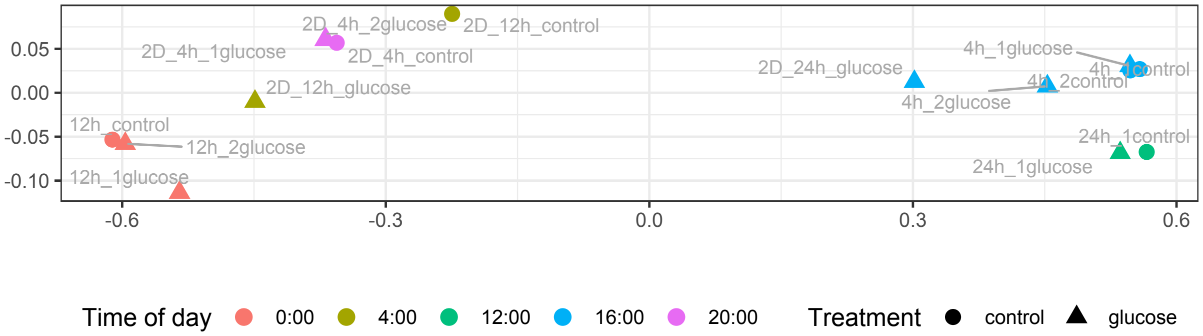


**Figure S11.** NMDS of *Prochlorococcus* metatranscriptomes. For each of the 15 samples the metatranscriptome consisted of the transcript levels ($log_2$, normalized) of the top 90% of genes. A 2-dimensional NMDS was performed on Euclidean distances calculated over 1 minus the Pearson correlation matrix for the sample metatranscriptomes. Stress was ~0. In the legend, the time of day indicates when the mRNA was fixed. Samples cluster primarily by day ($x$>0) and night ($x$<0), and secondarily by time of day regardless of glucose or control.

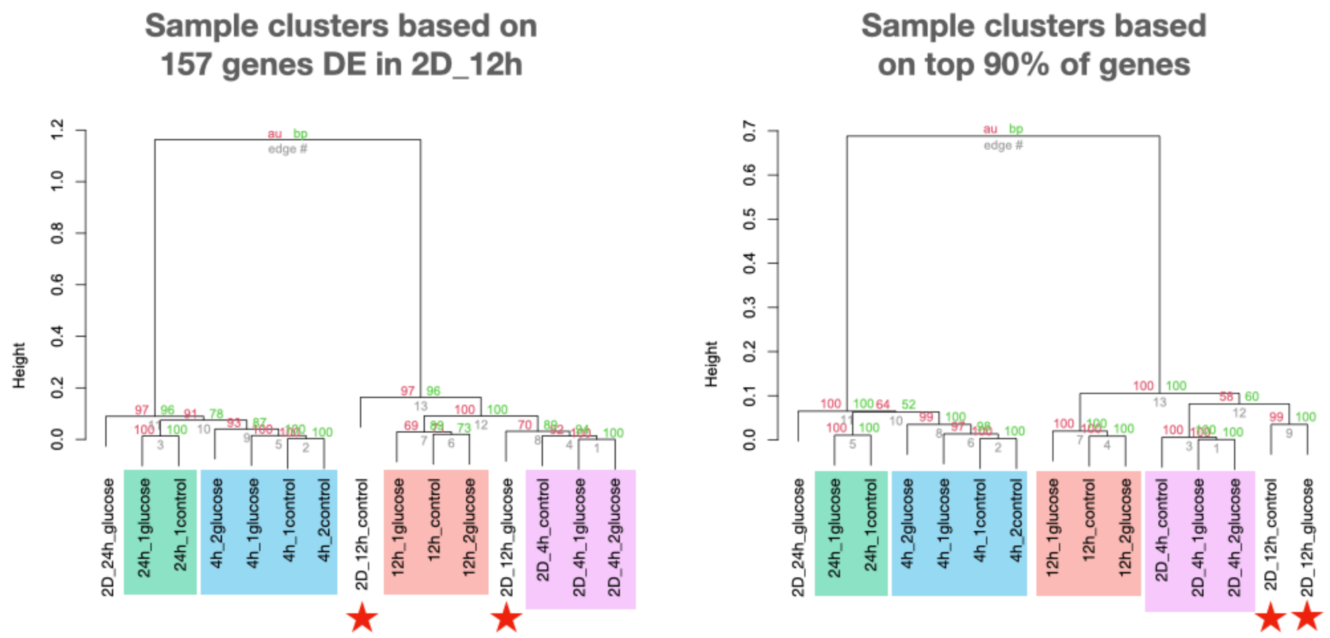


**Figure S12.** Sample dendrograms based on the genes that were differential expressed (DE) in 2D_12h glucose vs. control (left) and based on the top 90% of genes used in the NMDS analysis (right). For both dendrograms, correlation based Euclidean distances were calculated between samples, followed by Ward D2 clustering. 10,000 bootstraps were performed, and approximate unbiased *p*-values calculated for the clusters (red percentages, significant if >95) as well as traditional bootstrap support (green percentages), using the approach of Suzuki & Shimodaira (11) implemented in the R pvclust package. Clusters are shaded by collection time as in the NMDS Figure S11. As in the NMDS, samples cluster first by day vs. night and then by time of day. The exceptions are the 2D_12h samples (with red stars) in the left dendrogram, which suggests a disruption in diel expression patterns for the 157 that were DE in 2D_12h.

| Sample# | Date | Start | End | Light flux (E $m^{-2}$) |
|---|---|---|---|---|
| **Diel study** | | | | (per 4 hour) |
| 1 | 10/6/2017 | 0640 | 1040 | 5.0 |
| 2 | 10/6/2017 | 1245 | 1645 | 14.9 |
| 3 | 10/6/2017 | 1830 | 2235 | 0.0 |
| 4 | 10/7/2017 | 0035 | 0435 | 0.0 |
| 5 | 10/7/2017 | 0635 | 1035 | 10.7 |
| 6 | 10/7/2017 | 1245 | 1640 | 17.7 |
| 7 | 10/7/2017 | 1830 | 2235 | 0.0 |
| 8 | 10/8/2017 | 0030 | 0430 | 0.0 |
| 9 | 10/8/2017 | 0640 | 1040 | 12.3 |
| 10 | 10/8/2017 | 1240 | 1640 | 17.6 |
| **Genomics Exp. 1** | | | **Incubation time (h)** | **Total light (E $m^{-2}$)** |
| 1 | 10/6/17 | 1200 | 4 | 18.4 |
| 2 | 10/6/17 | 1200 | 12 | 20.9 |
| 3 | 10/6/17 | 1200 | 24 | 39.4 |
| **Genomics Exp. 2** | | | | |
| 1 | 10/7/2017 | 1600 | 4 | 3.3 |
| 2 | 10/7/2017 | 1600 | 12 | 3.3 |
| 3 | 10/7/2017 | 1600 | 24 | 40.4 |

**Table S1.** Date and sampling times for the diel and genomic incubation experiments and surface light flux during the incubations.

**Table S2**. List of 718 ASVs identified within the 16S rRNA data sets. This spreadsheet shows the ASV ID, total number of reads, scikit-learn annotation and sequence.

**Table S3**. List of 33 differentially abundant (DA) ASVs in response to glucose. The 33 DA ASVs always had fold changes that exceeded 1.2 and $p$-values <0.05.

**Table S4.** List of probes (6,834) and their sequences utilized in the arrays for 1,200 genes of different strains of *Prochlorococcus* that dominate at Station ALOHA.

**Table S5**. Detailed lists of normalized transcription values and description for the 775 genes in *Prochlorococcus.* Data spreadsheet shows the $\log_2$ transcript levels at each time point. This spreadsheet also shows the annotation, pathways of each gene, location, ecotype and COG function.

**Table S6.** Number of detected and undetected gene targets for all strains of *Prochlorococcus* represented on the microarray**.**

**Table S7.** List of 174 genes significantly differentially expressed (DE) in response to glucose. Expression changes for the 174 DE genes always exceeded 1.5-fold (mean 2.3-fold) and P value <0.05.

| **Pathway** | Genes included in gene set for EGSEA | Num HL | Num LL |
|---|---|---|---|
| Cell division | *ftsZ, minC, minD, minE* | 64 | 10 |
| Circadian rhythm | *kaiC* | 20 | - |
| Energy metab. | *atpA, atpB, atpC, atpE, atpF, atpI* | 51 | 8 |
| Photosystem I | *psaA, psaB* | 40 | 11 |
| Carbon fixation | *rbcL, rbcS, rbcS/L* | 25 | - |
| Respiration | *ctaC/coxB, catE/coxC, cyoB/coxA, cyoC* | 59 | 7 |
| Pentose phosphate | *opcA, rbsK, tal, zwf* | 74 | 5 |
| Glycolysis | *pykF, glk* | 39 | 2 |
| Glycolysis / Krebs cycle | *pdhA, pdhB, lpdA* | 36 | 11 |
| Krebs cycle and related genes | *fumC, ppc, sdhB* | 40 | 2 |
| Glycogen metab. | *glgC* | 20 | 2 |
| Glycolysis / gluconeogenesis | *eno, fda, pgi, cbbA, fda/cbbA* | 69 | 3 |
| Entner-Dudoroff | *eda, gdh* | 23 | 2 |
| Pyruvate metab. | *dld, ddA, lldD* | - | - |
| Sugar transporter | *glcH* | 16 | - |
| Amino acid metab. | *argH, glmS* | 32 | - |
| Nitrogen metab. | *glnA, glsF* | 35 | 2 |

**Table S8**. Differentially expressed pathways had collectively strong changes in the genes indicated. Pathways were separately defined for HL and LL *Prochlorococcus* and used all gene targets on the microarray for each gene symbol (Table S4). More HL than LL genes were detected for each pathway (rightmost two columns) likely because HL strains had more gene targets represented on the microarray and because we used surface water samples.

| | Glucose vs. Control | | | | | | | |
|---|---|---|---|---|---|---|---|---|
| | 1D 12h | | 1D 24h | | 2D 4h | | 2D 12h | |
| | HL Pro | LL Pro | HL Pro | LL Pro | HL Pro | LL Pro | HL Pro | LL Pro |
| Cell division | ↑ | o | o | o | o | o | o | o |
| Circadian rhythm | ↓ | | o | | ↑ | | o | |
| Energy metab. | o | o | o | o | o | o | o | o |
| Photosystem I | ↑ | o | ↓ | o | ↓ | ↓ | ↓ | o |
| Carbon fixation | o | | o | | o | | o | |
| Respiration | o | o | ↑ | o | o | o | ⬆ | o |
| Pentose phosphate | o | o | o | o | o | o | ⬆ | o |
| Glycolysis | o | o | o | o | o | o | ↓ | o |
| Glycolysis / Krebs cycle | o | o | o | o | o | o | o | o |
| Krebs cycle and related | ↓ | o | o | o | o | o | o | o |
| Glycogen metab. | o | o | o | o | o | o | ↓ | o |
| Glycolysis / gluconeogenesis | o | o | o | o | o | o | o | o |
| Entner-Dudoroff | o | o | o | o | o | o | o | o |
| Pyruvate metab. | | | | | | | | |
| Sugar transporter | o | | o | | ↑ | | ⬆ | |
| Amino acid metab. | ↓ | | o | | ↑ | | o | |
| Nitrogen metab. | o | o | ↓ | o | o | o | o | o |

**Table S9**. Summarize of the EGSEA results for differentially expressed pathways for HL and LL ecotypes. Red and blue arrows indicate significant ($p < 0.01$) responses to glucose. Thick arrows indicate that the average fold change was ≥1.5×. ° means the change was not significant. Empty cells mean that genes in the pathway were not detected.

**Table S10.** List of the EGSEA results for differentially expressed pathways for HL and LL ecotypes. Adjusted *p*-values < 0.01 were significant.
